# Multivalent BBSome – IFT proximities support IFT-A – IFT-B coordination *in vivo*

**DOI:** 10.64898/2026.11.11.765876

**Authors:** Zijie Shen, Guanghan Chen, Ming Li, Yongping Chai, Wei Li, Guangshuo Ou

## Abstract

Intraflagellar transport (IFT) depends on coordinated movement of IFT-A and IFT-B, yet how these assemblies remain coupled within dynamic ciliary trains is unclear. Here, we combine endogenous fluorescent knock-in tagging, quantitative *in vivo* FLIM-FRET, structural modeling, and targeted mutagenesis to define IFT organization in *C. elegans* and mammalian cells. FLIM-FRET mapping identifies proximity networks within IFT-B and places IFT144 near IFT172 and IFT88 in intact cilia. Loss of BBSome function is associated with weaker IFT-A – IFT-B proximity signals and divergent subcomplex transport dynamics, consistent with a contribution of the BBSome to IFT coordination *in vivo*. Split-GFP complementation places BBSome components near IFT-A and IFT-B, while structural modeling nominates complementary charged surfaces on the IFT46 – IFT52 dimer and BBS7/BBS9. Charge-reversal mutations targeting these surfaces impair BBSome recruitment and functional rescue. Together, these findings support a model in which multivalent BBSome – IFT proximities contribute to IFT-A – IFT-B coordination during ciliary transport.

## Introduction

Cilia are evolutionarily conserved, microtubule-based organelles that function as essential signaling hubs to regulate different physiological processes including sensory perception and developmental patterning [1][2][3][4][5]. Cilia rely on intraflagellar transport (IFT) to assemble, maintain and remodel their signaling compartments [6][7][8][9]. IFT is mediated by large protein trains that move along axonemal microtubules and coordinate the trafficking of structural and signaling cargoes [10][11]. These trains contain two major subcomplexes, IFT-A and IFT-B, which have distinct molecular compositions and trafficking functions but move together during processive transport [12][13][14]. How structurally defined IFT-A – IFT-B interfaces are maintained and regulated within dynamic trains in living cilia remains incompletely understood.

Recent structural studies have provided high-resolution views of IFT-A and IFT-B complexes and have begun to define their organization within assembled IFT trains [15][16][17][18][19][20][21]. These studies revealed a modular architecture for the IFT machinery and suggested potential points of contact between IFT-A and IFT-B. However, several candidate interface regions remain flexible, poorly resolved or absent from current structural maps [22][23][24]. Moreover, static structural models cannot fully explain how IFT-A and IFT-B maintain coordinated movement during continuous train assembly, remodeling and bidirectional transport *in vivo*. Direct measurements of subunit proximity within intact cilia are therefore needed to define how IFT trains are organized under native cellular conditions.

The BBSome is a conserved ciliary trafficking complex best known for mediating the transport and recycling of membrane-associated ciliary cargoes [25][26][27][28][29]. Loss of BBSome function causes Bardet – Biedl syndrome (BBS) and disrupts ciliary signaling, cargo distribution, and IFT dynamics [30][31][32]. Genetic and imaging studies in *C. elegans* and vertebrate systems have suggested that the BBSome also contributes to IFT-train organization, because IFT-A and IFT-B display altered localization and transport behavior when BBSome function is impaired [32][33]. However, whether the BBSome stabilizes physical coupling between IFT-A and IFT-B, and what molecular features underlie this regulation, remain unknown.

Here we investigate BBSome-associated coordination of IFT-A and IFT-B *in vivo*. Using endogenous fluorescent knock-in strains in *C. elegans*, we establish a native platform for quantitative IFT imaging and FLIM-FRET-based proximity mapping. This approach identifies proximity relationships within IFT-B and reveals BBSome-dependent proximity between IFT144 and the IFT-B subunits IFT172 and IFT88. Parallel experiments in mammalian cells are consistent with a similar contribution of the BBSome to IFT-A – IFT-B coordination. We further place the BBSome in proximity to both IFT-A- and IFT-B-associated sites and identify candidate complementary charged surfaces on the acidic IFT46 – IFT52 module and BBS7 or BBS9. Charge-reversal mutations targeting these surfaces reduce BBSome recruitment and ciliary function. Together, our findings support a working model in which the BBSome contributes to IFT train coordination through multivalent proximity relationships during ciliary transport.

## Results

### Endogenous fluorescent tagging establishes a native platform for quantitative analysis of IFT dynamics *in vivo*

To define the organization of IFT trains under native expression conditions, we generated a panel of *C. elegans* knock-in strains in which endogenous IFT-particle subunits were tagged at their C termini with GFP or mScarlet. This collection includes all six IFT-A components and fourteen IFT-B components encoded in *C. elegans* (Fig. 1A and Fig. S1). Because the *C. elegans* IFT-B complex lacks IFT25 and IFT27, which are present in several other ciliated organisms [12], this panel represents a near-complete endogenous fluorescent atlas of the worm IFT machinery.

**Figure 1.**
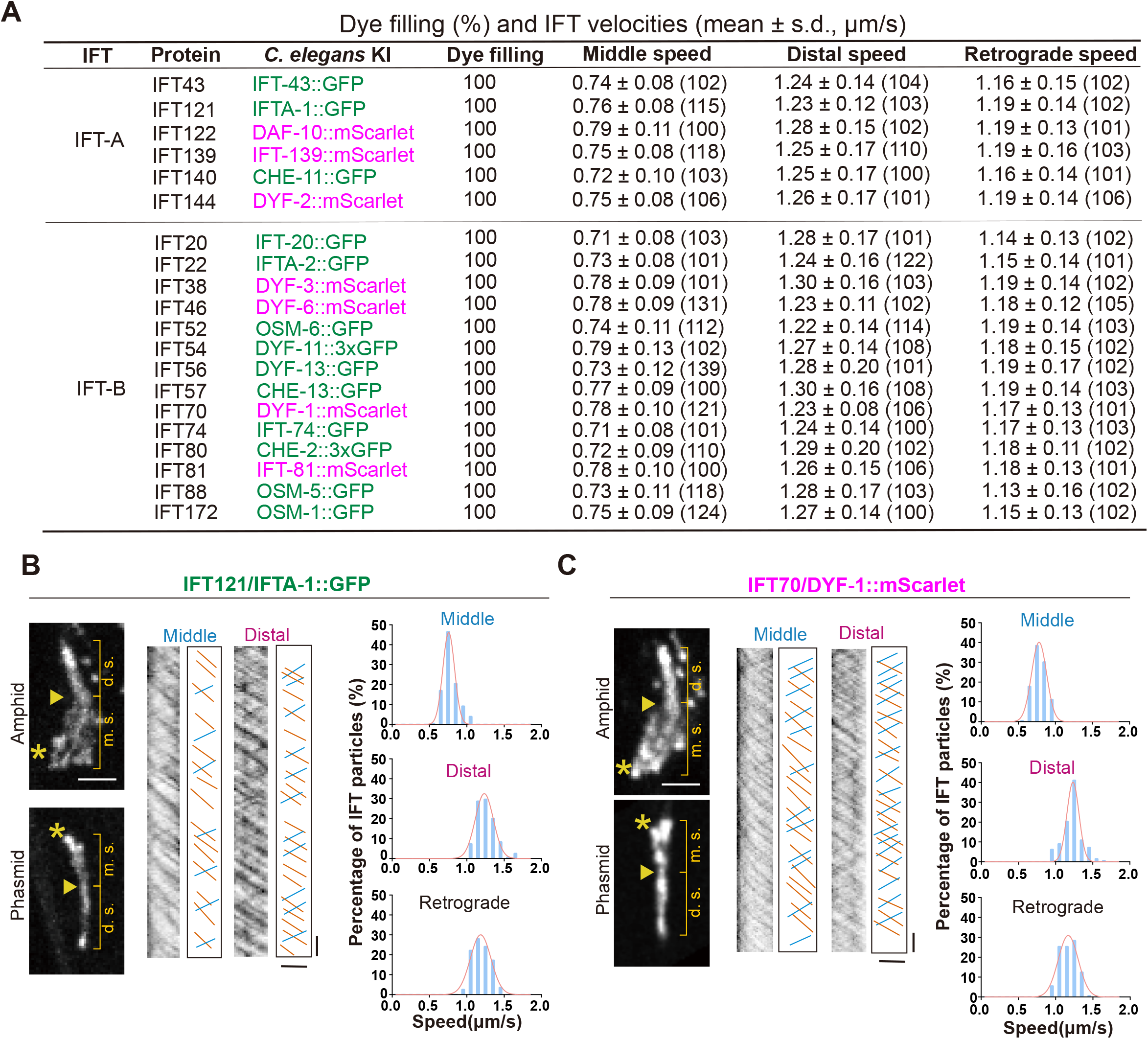
Endogenous fluorescent knock-in strains reveal coordinated transport dynamics of IFT-A and IFT-B subunits in *C. elegans*. **(A)** Summary of endogenous GFP- or mScarlet-tagged knock-in strains for *C. elegans* IFT-A and IFT-B subunits, together with dye-filling percentages and measured IFT velocities in amphid and phasmid sensory cilia. Middle and distal segment velocities correspond to anterograde transport; retrograde velocities were measured along the ciliary shaft. Velocity data are mean ± s.d.; numbers in parentheses indicate the number of transport events analyzed. Data are from at least 10 independent phasmid cilia of *C. elegans*. Dye filling was scored in more than 100 animals per strain. **(B)** Representative ciliary localization, kymographs, and velocity distributions of the IFT-A subunit IFT121/IFTA-1::GFP. Left, representative amphid and phasmid cilia. Asterisks mark the ciliary base/transition zone, and arrowheads mark the junction between the middle and distal segments. Scale bar, 2.5 μm. Middle, representative kymographs showing IFT transport in the middle and distal segments; orange lines indicate anterograde IFT tracks, and blue lines indicate retrograde IFT tracks. Horizontal scale bar, 2.5 μm; vertical scale bar, 2.5 s. Right, velocity distributions and Gaussian fits for IFT121/IFTA-1::GFP transport in the middle segment, distal segment, and retrograde direction. **(C)** Representative ciliary localization, kymographs, and velocity distributions of the IFT-B subunit IFT70/DYF-1::mScarlet, displayed as in (B).

Live imaging of amphid and phasmid sensory cilia revealed that all tagged IFT-A and IFT-B components localized to cilia and underwent robust bidirectional transport. Across the knock-in panel, anterograde transport showed the expected segment-specific kinetics, with particles moving at approximately 0.7 μm/s in the middle segment and approximately 1.2 – 1.3 μm/s in the distal segment, whereas retrograde transport occurred at approximately 1.1 – 1.2 μm/s (Fig. 1A – C and Fig. S1). These velocities are consistent with previously reported IFT dynamics in *C. elegans* sensory cilia [33], supporting the fidelity of the endogenous tagging strategy.

Importantly, IFT-A and IFT-B components displayed highly similar transport behaviors across the ciliary shaft, indicating that the two subcomplexes move coordinately within intact IFT trains under basal conditions. The knock-in strains also retained normal sensory cilium function, as assessed by DiI dye-filling assays in more than 100 animals per strain (Fig. 1A). Thus, C-terminal fluorescent tagging did not detectably perturb IFT localization, transport dynamics, or sensory cilium function under the conditions tested. Together, these knock-in lines establish a native, quantitative platform for comparing IFT-particle dynamics and for probing the molecular organization of IFT trains *in vivo*.

### *In vivo* FLIM-FRET maps proximity relationships within the IFT-B complex

To probe the molecular organization of IFT trains in living cilia, we next applied fluorescence lifetime imaging microscopy – Förster resonance energy transfer (FLIM-FRET) [34][35] to the endogenous knock-in strains. In this assay, energy transfer from GFP-tagged donor subunits to mScarlet-tagged acceptor subunits reduces the GFP fluorescence lifetime, providing a quantitative readout of molecular proximity within intact cilia. We therefore generated double knock-in strains by genetic crossing and measured GFP lifetimes directly in sensory cilia under native expression conditions.

As an initial validation of this strategy, we examined IFT-B subunit pairs whose C-termini are positioned in close proximity in the available IFT-B structural model [18][22]. Significant reductions in GFP lifetime were detected for IFT57/CHE-13::GFP with IFT38/DYF-3::mScarlet, IFT80/CHE-2::GFP with IFT54/DYF-11::mScarlet, IFT74::GFP with IFT81::mScarlet, and IFT52/OSM-6::GFP with IFT46/DYF-6::mScarlet (Fig. 2). These lifetime shifts are consistent with the predicted spatial arrangement of the corresponding C-termini and indicate that endogenous FLIM-FRET can resolve subunit proximities within native IFT trains.

**Figure 2.**
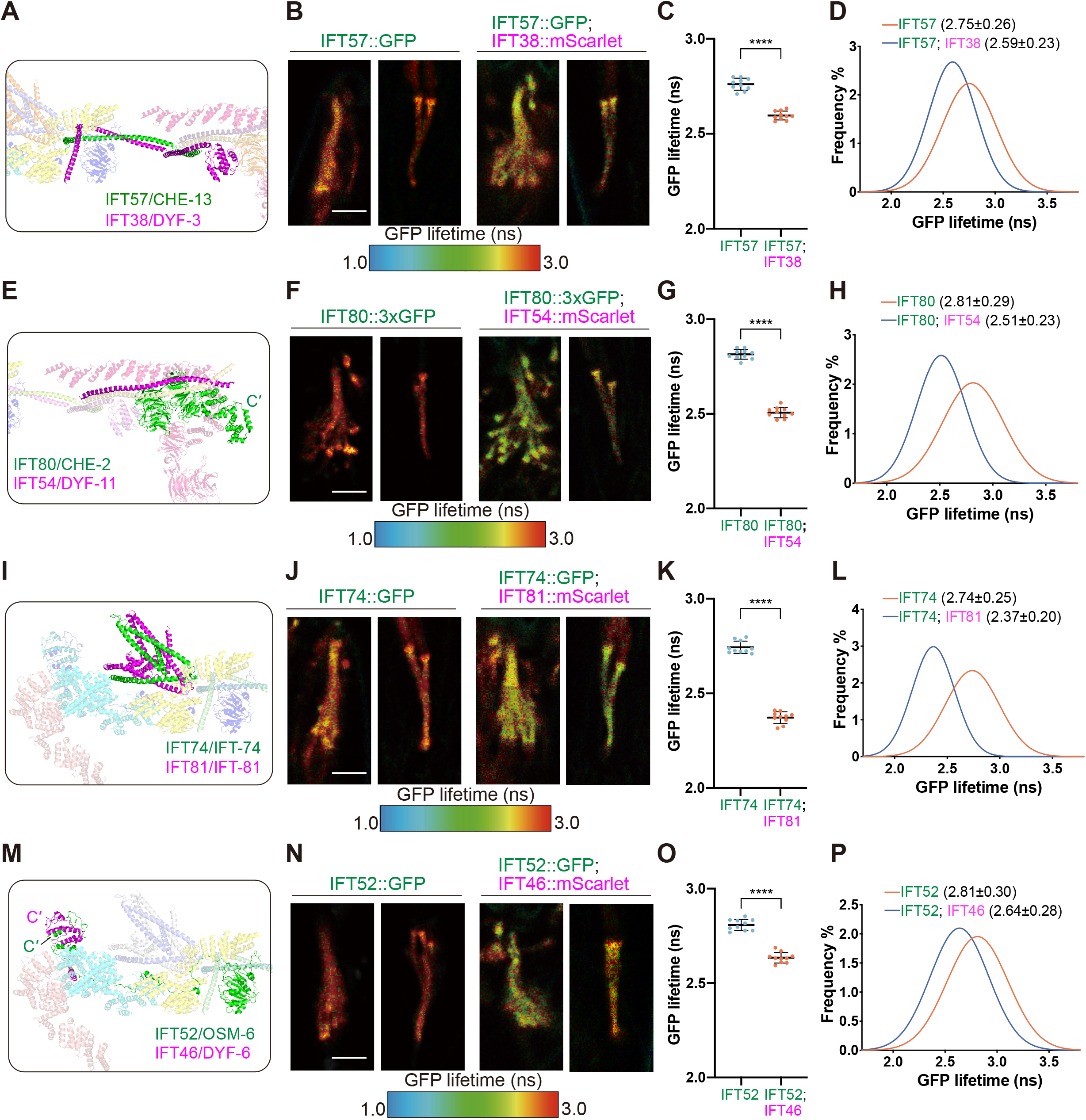
Endogenous FLIM-FRET identifies proximity relationships within the IFT-B complex in vivo. (A, E, I, and **M)** Structural models of the IFT-B complex showing the relative positions of the indicated subunit pairs, including IFT57/CHE-13 and IFT38/DYF-3 (A), IFT80/CHE-2 and IFT54/DYF-11 (E), IFT74 and IFT81 (I), and IFT52/OSM-6 and IFT46/DYF-6 (M). C′ denotes the C-terminal end. **(B, F, J, and N)** Representative FLIM images of cilia expressing the indicated GFP-tagged donor alone or together with the corresponding mScarlet-tagged acceptor. GFP fluorescence lifetime is displayed as a pseudocolor map from 1.0 to 3.0 ns. Scale bars, 2.5 μm. **(C, G, K, and O)** Quantification of GFP lifetimes from the corresponding FLIM images. Each dot represents the apparent mean GFP lifetime of one cilium, calculated by fitting the fluorescence decay curve of that cilium with a mono-exponential decay model. N = 10 cilia from 10 animals per genotype. Bars show mean ± s.d. ****P < 0.0001, two-tailed Student’s t-test. **(D, H, L, and P)** GFP fluorescence lifetime distributions and Gaussian fits for the corresponding donor-only and donor–acceptor strains. Curves show distributions from the same 10 ciliary regions of interest per genotype. Values in parentheses indicate the mean ± s.d. of the cilium-level apparent mean lifetimes obtained from Gaussian fits.

To assess assay specificity, we analyzed selected comparison pairs, including IFT140/CHE-11 with IFT144/DYF-2, IFT172/OSM-1 with IFT38/DYF-3, and IFT140/CHE-11 with IFT38/DYF-3. None showed a significant GFP lifetime reduction under the conditions tested (Fig. S2A – L). Detectable lifetime shifts were observed for selected structurally supported pairs, whereas no shift was detected for these comparison pairs (Fig. S2M). These results support the use of endogenous FLIM-FRET as an *in vivo* proximity readout for the tested pairs. FLIM-FRET reports fluorophore proximity rather than binding affinity, and a negative result does not exclude proximity because fluorophore separation, dipole orientation, and donor – acceptor occupancy influence FRET.

### BBSome loss is associated with reduced IFT144 – IFT-B FLIM-FRET and divergent IFT-A – IFT-B transport

Having established endogenous FLIM-FRET as an *in vivo* proximity assay, we next sought to identify proximity relationships that could report on IFT-A – IFT-B coupling within intact trains. Available *in situ* IFT-train models place the IFT-A subunit IFT144/DYF-2 near the IFT-B subunits IFT172/OSM-1 and IFT88/OSM-5 (Fig. 3A) [22]. Biochemical mapping has identified an IFT-A – IFT-B interface involving IFT122 – IFT144 and IFT52 – IFT88 [36], whereas current train structures position the flexible C terminus of IFT172 near the C-terminal region of IFT144 [22]. However, the corresponding C-terminal regions are flexible or poorly resolved in current structural maps, leaving the local organization of this putative IFT-A – IFT-B contact unclear. We therefore used AlphaFold 3 modeling of the *C. elegans* C-terminal regions of IFT144/DYF-2, IFT172/OSM-1, and IFT88/OSM-5 to nominate a candidate proximity relationship (Fig. 3B). Similar configurations were predicted for the corresponding human and *Chlamydomonas* proteins, suggesting that this region may represent a conserved IFT-A – IFT-B proximity site (Fig. S3A – C).

**Figure 3.**
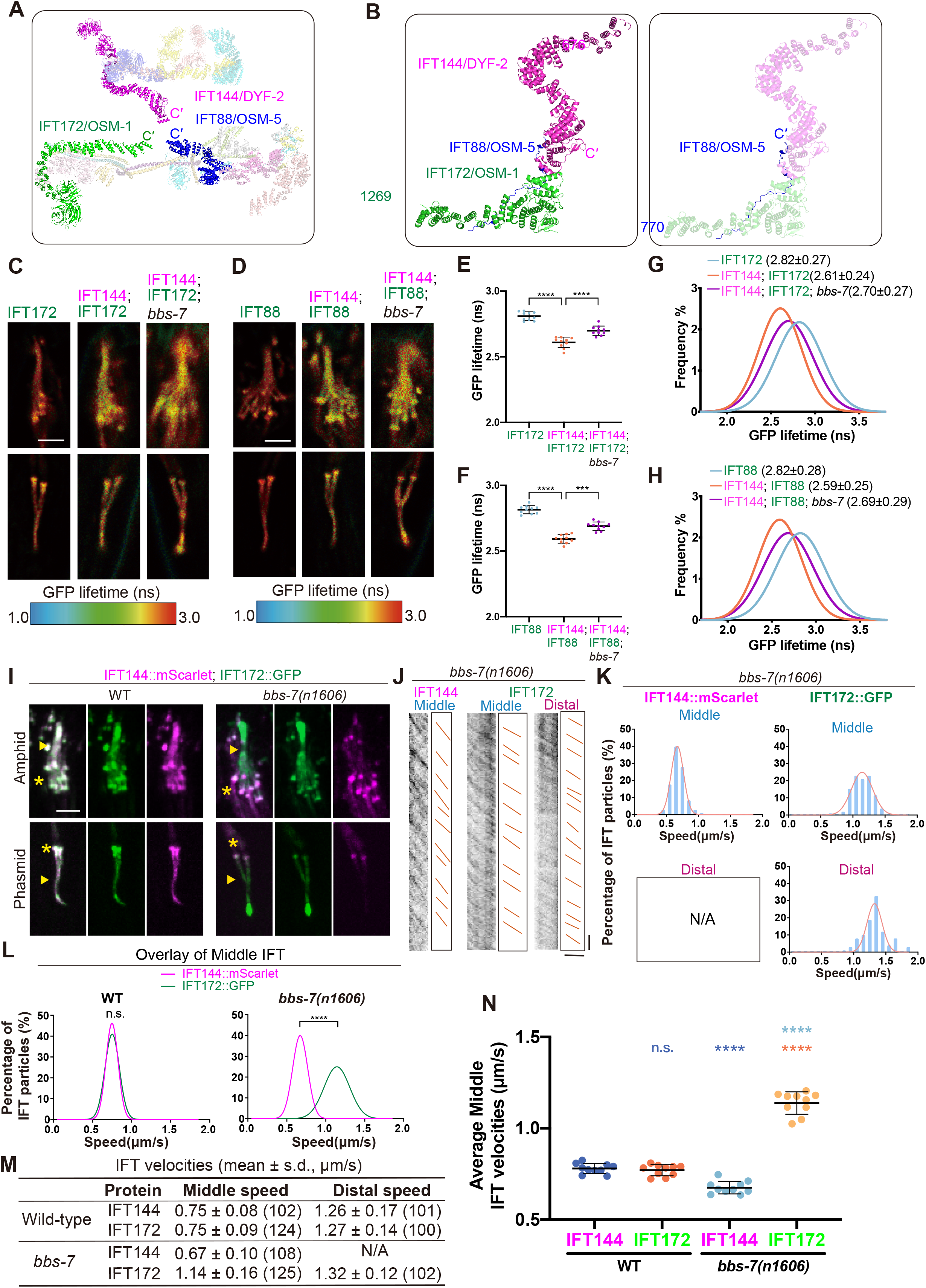
BBSome loss is associated with reduced IFT144–IFT-B FLIM-FRET and divergent transport in *C.elegans*. **(A)** Model of the anterograde IFT train based on available IFT-A (8BDA) and IFT-B (8BD7) structural models, showing the predicted proximity of IFT144/DYF-2 to the IFT-B subunits IFT172/OSM-1 and IFT88/OSM-5. C′ denotes the C-terminal end. **(B)** AlphaFold 3-predicted model of the C-terminal regions of *C. elegans* IFT144/DYF-2, IFT172/OSM-1, and IFT88/OSM-5. The left panel highlights IFT144/DYF-2 residues 676–1,364 and IFT172/OSM-1 residues 1,269–1,737, whereas the right panel highlights IFT88/OSM-5 residues 770–820. C′ denotes the C-terminal end. **(C)** Representative FLIM images of cilia expressing IFT172/OSM-1::GFP alone, IFT172/OSM-1::GFP together with IFT144/DYF-2::mScarlet in WT animals, or IFT172/OSM-1::GFP together with IFT144/DYF-2::mScarlet in *bbs-7(n1606)* animals. GFP fluorescence lifetime is displayed as a pseudocolor map from 1.0 to 3.0 ns. Scale bar, 2.5 μm. **(D)** Representative FLIM images of cilia expressing IFT88/OSM-5::GFP alone, IFT88/OSM-5::GFP together with IFT144/DYF-2::mScarlet in WT animals, or IFT88/OSM-5::GFP together with IFT144/DYF-2::mScarlet in *bbs-7(n1606)* animals. GFP fluorescence lifetime is displayed as in (C). Scale bar, 2.5 μm. **(E and F)** Quantification of GFP lifetimes for IFT172/OSM-1::GFP (E) and IFT88/OSM-5::GFP (F) from cilia shown in (C) and (D), respectively. Each dot represents the apparent mean GFP lifetime of one cilium, calculated by fitting the fluorescence decay curve of that cilium with a mono-exponential decay model. N = 10 cilia from 10 animals per genotype. Bars show mean ± s.d. ***P < 0.001; ****P < 0.0001, P values were calculated using one-way ANOVA followed by Tukey’s multiple-comparisons test. **(G and H)** GFP fluorescence lifetime distributions and Gaussian fits for IFT172/OSM-1::GFP (G) and IFT88/OSM-5::GFP (H) in the strains shown in (C) and (D), respectively. Curves show distributions from the same 10 ciliary regions of interest per genotype. Values in parentheses indicate the mean ± s.d. of the cilium-level apparent mean lifetimes obtained from Gaussian fits. **(I)** Representative ciliary images of endogenous IFT144/DYF-2::mScarlet and IFT172/OSM-1::GFP in WT and *bbs-7(n1606)* animals. Asterisks mark the ciliary base/transition zone, and arrowheads mark the junction between the middle and distal segments. Scale bar, 2.5 μm. **(J)** Representative kymographs showing anterograde transport of IFT144/DYF-2::mScarlet and IFT172/OSM-1::GFP in *bbs-7(n1606)* cilia. Orange lines indicate anterograde IFT tracks. Horizontal scale bar, 2.5 μm; vertical scale bar, 2.5 s. **(K)** Velocity distributions and Gaussian fits for anterograde IFT172/OSM-1::GFP and IFT144/DYF-2::mScarlet transport in *bbs-7(n1606)* animals. N/A, not applicable. **(L)** Overlay plots of middle-segment velocities of IFT144/DYF-2::mScarlet and IFT172/OSM-1::GFP in WT and *bbs-7(n1606)* animals. Event-level distributions are shown descriptively; cilium-level inferential statistics are reported in (N). n.s., not significant; ****P < 0.0001, two-tailed Student’s t-test. **(M)** Summary of anterograde IFT velocities for IFT144/DYF-2::mScarlet and IFT172/OSM-1::GFP in wild-type and *bbs-7(n1606)* animals. Data are mean ± s.d.; numbers in parentheses indicate the number of transport events analyzed. N/A, not applicable. Data are from at least 10 independent phasmid cilia of *C. elegans*. **(N)** Average middle IFT velocities of IFT144 and IFT172 from 10 independent cilia. n.s., not significant; ****P < 0.0001, P values were calculated using one-way ANOVA followed by Tukey’s multiple-comparisons test.

Consistent with this model, endogenous FLIM-FRET revealed a significant reduction in GFP lifetime when IFT172/OSM-1::GFP or IFT88/OSM-5::GFP was paired with IFT144/DYF-2::mScarlet in wild-type *C. elegans* cilia (Fig. 3C – H). These data indicate that IFT144 lies in close proximity to both IFT172 and IFT88 within native IFT trains, providing a molecular readout for IFT-A – IFT-B organization *in vivo*.

We then asked whether this IFT144 – IFT-B proximity depends on BBSome function. Previous work suggested that BBSome dysfunction separates IFT-A and IFT-B transport in *C. elegans* sensory cilia [33], but whether this reflects altered molecular proximity between the two subcomplexes under native expression conditions has remained unclear. To test this, we introduced endogenous IFT144/DYF-2::mScarlet together with IFT172/OSM-1::GFP or IFT88/OSM-5::GFP into the *bbs-7(n1606)* mutant background.

In wild-type animals, IFT144/DYF-2 and IFT172/OSM-1 displayed similar anterograde velocities in the middle segment. In contrast, in *bbs-7* mutants, IFT144/DYF-2::mScarlet was largely restricted to the middle segment and moved at approximately 0.67 μm/s, whereas IFT172/OSM-1::GFP entered the distal segment and moved significantly faster, at approximately 1.14 μm/s in the middle segment and 1.32 μm/s in the distal segment (Fig. 3I – N). Mutants of *bbs-1*, *bbs-2*, *bbs-8* and *bbs-9* showed similar IFT144 and IFT172 localization defects (Fig. S3D), and BBS-9::GFP failed to enter cilia in *bbs-7* mutants (Fig. S3E), indicating broader disruption of BBSome function in the *bbs-7* background. We therefore used *bbs-7(n1606)* as a representative BBSome-deficient condition for subsequent analyses.

In *bbs-7* mutant worms, whole-cilium FLIM-FRET lifetime shifts between IFT144/DYF-2::mScarlet and either IFT172/OSM-1::GFP or IFT88/OSM-5::GFP were reduced, with donor lifetimes shifting toward donor-only values (Fig. 3C – H). The reduced lifetime shift is therefore consistent with altered IFT-A – IFT-B proximity or co-occupancy.

To ask whether BBSome-associated IFT coordination extends to mammalian cilia, we generated a BBS7-knockout IMCD3 cell line and stably expressed fluorescently tagged IFT144 and IFT88 by lentiviral transduction. The knockout cells showed a canonical BBSome-deficiency phenotype, including ciliary accumulation of Smoothened (SMO) [37], confirming disruption of BBS7-dependent ciliary trafficking (Fig. 4A,B). In contrast to *C. elegans bbs-7* mutants, both IFT144 and IFT88 entered the full length of IMCD3 cilia in BBS7-knockout cells (Fig. 4C). However, live-cell imaging revealed that their transport dynamics were no longer matched. For transport measurements, we transiently expressed IFT144::3xNeoGreen and IFT88::3xNeoGreen under the same imaging conditions. IFT144 and IFT88 moved at similar anterograde velocities in wild-type cells, whereas BBS7 loss altered the anterograde velocities of both reporters, with IFT88 moving significantly faster than IFT144 in the knockout background (Fig. 4D – H). Thus, at the population level, BBS7 loss is associated with divergent IFT-A and IFT-B transport velocities in mammalian cilia even when both subcomplexes retain ciliary access.

**Figure 4.**
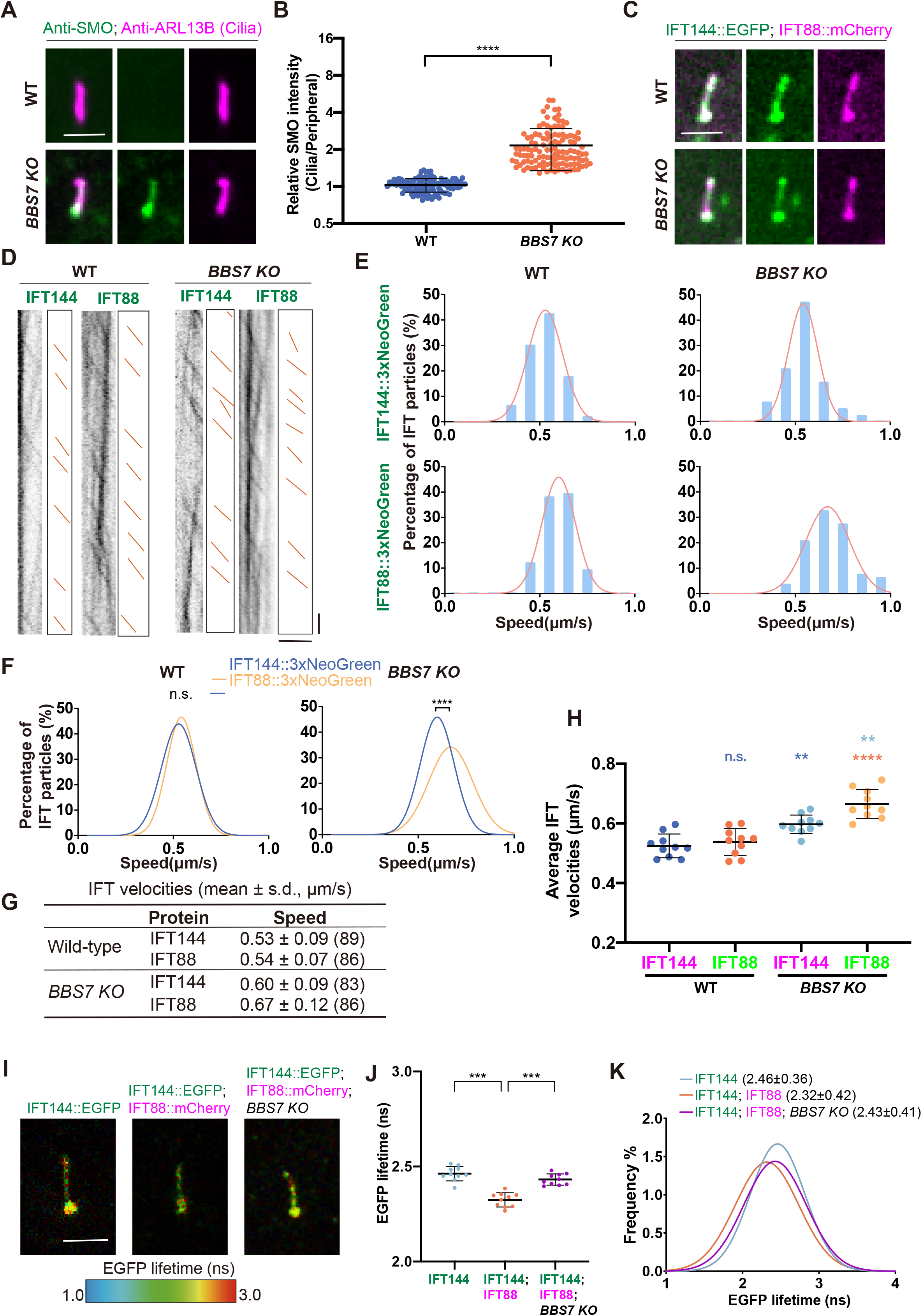
BBS7 loss is associated with divergent IFT144 and IFT88 transport and reduced FLIM-FRET in mammalian cilia. (A) Representative immunofluorescence images of SMO in WT and *BBS7 KO* cilia. ARL13B was used as a ciliary marker. Scale bar, 2.5 μm. **(B)** Quantification of relative SMO enrichment in WT and *BBS7 KO* cilia, calculated as the ratio of ciliary to peripheral SMO signal intensity. More than 100 cilia from at least three independent experiments were analyzed per condition. ****P < 0.0001, two-tailed Student’s t-test. The y-axis is on a log2 scale. **(C)** Representative ciliary images of WT and *BBS7 KO* IMCD3 cells expressing IFT144::EGFP and IFT88::mCherry by lentiviral transduction. Scale bar, 2.5 μm. **(D)** Representative kymographs showing anterograde IFT transport of IFT144::3xNeoGreen and IFT88::3xNeoGreen in WT and *BBS7 KO* IMCD3 cells after transient transfection in separate experiments. Orange lines indicate anterograde IFT tracks. Horizontal scale bar, 2.5 μm; vertical scale bar, 2.5 s. **(E)** Velocity distributions and Gaussian fits for anterograde IFT144::3xNeoGreen and IFT88::3xNeoGreen transport in WT and *BBS7 KO* IMCD3 cells. **(F)** Overlay plots of IFT144::3xNeoGreen and IFT88::3xNeoGreen velocities in WT and *BBS7 KO* backgrounds. Event-level distributions are shown descriptively; cilium-level inferential statistics are reported in (H). n.s., not significant; ****P < 0.0001, two-tailed Student’s t-test. **(G)** Summary of anterograde IFT velocities for IFT144::3xNeoGreen and IFT88::3xNeoGreen in WT and *BBS7 KO* IMCD3 cells. Data are mean ± s.d.; numbers in parentheses indicate the number of transport events analyzed. Each group of IFT velocities was measured from at least 10 cells in separate IFT144 and IFT88 experiments. **(H)** Average IFT velocities of IFT144 and IFT88 from 10 independent cilia in 10 cells. n.s., not significant; **P < 0.01; ****P < 0.0001, P values were calculated using one-way ANOVA followed by Tukey’s multiple-comparisons test. **(I)** Representative FLIM images of IMCD3 cilia expressing IFT144::EGFP alone, IFT144::EGFP together with IFT88::mCherry in WT cells, or IFT144::EGFP together with IFT88::mCherry in *BBS7 KO* cells by lentiviral transduction. EGFP fluorescence lifetime is displayed as a pseudocolor map from 1.0 to 3.0 ns. Scale bar, 2.5 μm. **(J)** Quantification of EGFP lifetimes from cilia shown in (I). Each dot represents the apparent mean lifetime of one cilium. N = 10 cilia from 10 cells per condition. Bars show mean ± s.d. ***P < 0.001, P values were calculated using one-way ANOVA followed by Tukey’s multiple-comparisons test. **(K)** EGFP fluorescence lifetime distributions and Gaussian fits from cilia shown in (I). Curves show distributions from the same 10 ciliary regions of interest per condition. Values in parentheses indicate the mean ± s.d. of the cilium-level apparent mean lifetimes obtained from Gaussian fits.

We next used FLIM-FRET to determine whether this transport uncoupling was accompanied by altered molecular proximity between IFT144 and IFT88. In wild-type IMCD3 cells, IFT144::GFP showed a reduced fluorescence lifetime in the presence of IFT88::mCherry, consistent with close proximity between these subunits within mammalian IFT trains. In *BBS7*-knockout cells, the IFT144::GFP lifetime shifted back toward the donor-only baseline, consistent with reduced IFT144 – IFT88 proximity (Fig. 4I – K). Together with the endogenous knock-in analysis in *C. elegans*, these findings extend the association to a mammalian cell model.

### Multivalent BBSome – IFT proximities motivate a working model for IFT-A – IFT-B coordination

The reduced IFT144 – IFT172 and IFT144 – IFT88 FLIM-FRET signals in *bbs- 7* mutants were associated with altered IFT-A – IFT-B organization. We therefore asked whether the BBSome might contribute to this organization through proximity to multiple sites on the IFT machinery. Endogenous BBS-7::GFP localized to sensory cilia and displayed segment-specific transport dynamics comparable to those of IFT-A and IFT-B subunits, moving at approximately 0.75 μm/s in the middle segment, 1.21 μm/s in the distal segment, and 1.16 μm/s during retrograde transport (Fig. S4A – D).

These observations are consistent with the BBSome moving with IFT trains under basal conditions.

Although the precise position of the BBSome within anterograde IFT trains remains unresolved, available IFT-train models indicate that major surfaces of the IFT-A – IFT-B scaffold are occupied by kinesin-II, the ciliary membrane, or dynein-2 [24]. We therefore considered a peripheral BBSome position near the IFT-B scaffold and in potential proximity to IFT-A as a working model (Fig. 5A). To identify candidate BBSome – IFT proximity sites for experimental testing, we aligned predicted *C. elegans* BBSome models to reported BBSome structures [38][39][40] and examined their potential proximity to IFT subunits in AlphaFold 3 – based models. We did not obtain confident BBSome contacts with the IFT144/DYF-2 – IFT172/OSM-1 – IFT88/OSM-5 region analyzed above, suggesting that this FLIM-FRET reporter site reflects BBSome-dependent IFT-A – IFT-B organization rather than a direct BBSome docking site. Instead, the models suggested potential BBSome proximity to the C-terminal regions of the IFT-B subunits IFT46/DYF-6 and IFT52/OSM-6, as well as to the N-terminal region of IFT43 (Fig. 5B,C). These models were used as guides for experimental testing rather than as standalone evidence for binding.

**Figure 5.**
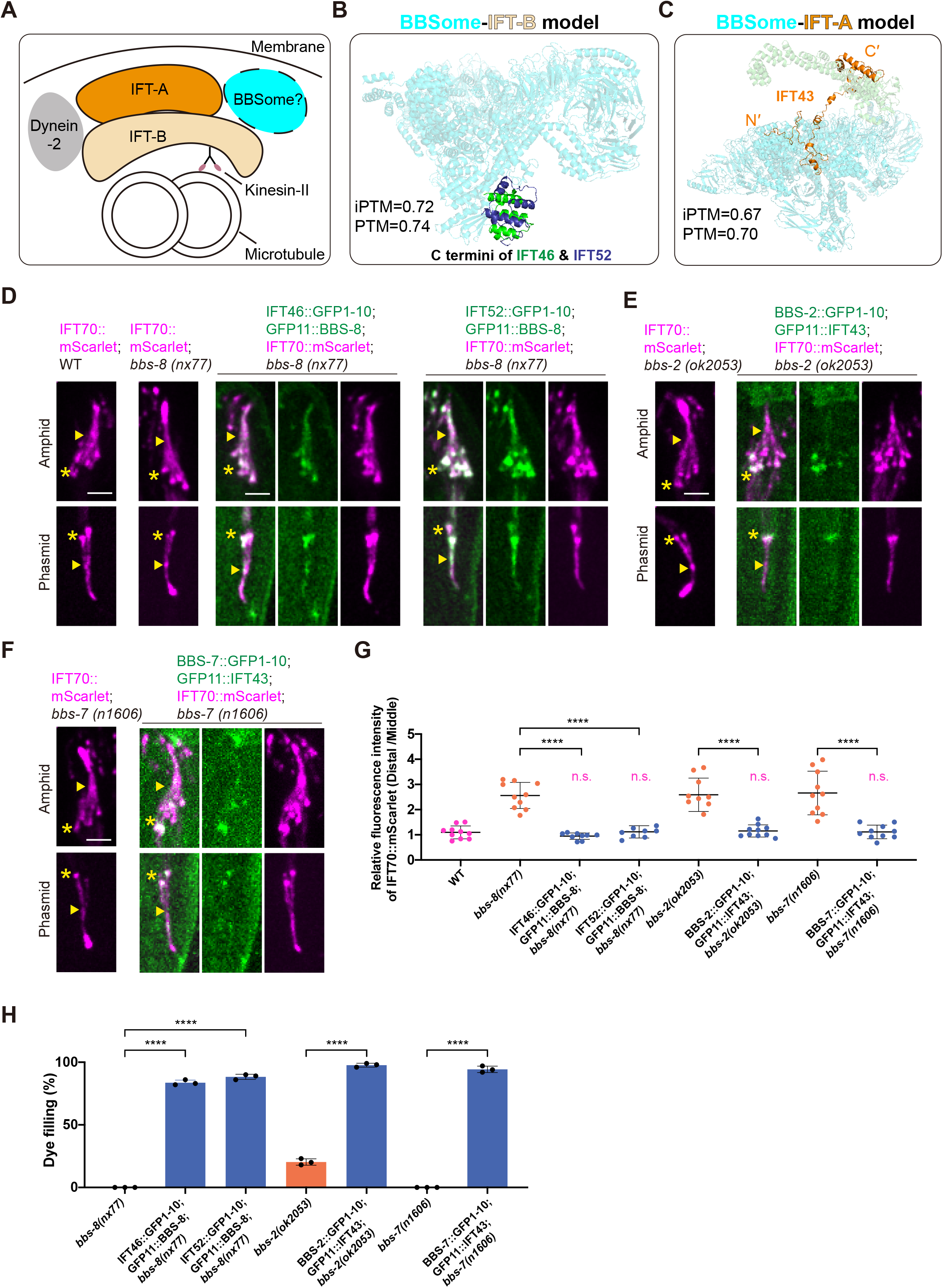
Multivalent BBSome–IFT proximities motivate a working model for IFT-A–IFT-B coordination. (A) Working model illustrating a potential position of the BBSome near the periphery of an anterograde IFT train, in proximity to both IFT-A and IFT-B. The precise position of the BBSome within intact IFT trains remains unresolved. **(B)** AlphaFold 3-based model of the *C. elegans* BBSome with the C-terminal regions of the IFT-B subunits IFT46/DYF-6 and IFT52/OSM-6. Shown are IFT46/DYF-6 residues 368–446 and IFT52/OSM-6 residues 369–427. The predicted *C. elegans* BBSome model was aligned to the reported bovine BBSome structure (6VNW). **(C)** AlphaFold 3-based model of the *C. elegans* BBSome with full-length IFT43. IFT121 was included in the model to constrain the IFT-A-associated configuration. The predicted *C. elegans* BBSome model was aligned to the reported bovine BBSome structure (6VNW). N′ and C′ denote the N- and C-terminal ends of IFT43, respectively. **(D)** Representative ciliary images showing split-GFP complementation between BBS-8 and the IFT-B subunits IFT46/DYF-6 or IFT52/OSM-6 in *bbs-8(nx77)* animals. IFT70/DYF-1::mScarlet was co-expressed as a ciliary IFT marker. WT and *bbs-8(nx77)* animals expressing IFT70/DYF-1::mScarlet alone are shown as controls. Asterisks mark the ciliary base/transition zone, and arrowheads mark the junction between the middle and distal segments. Scale bars, 2.5 μm. **(E)** Representative ciliary images showing split-GFP complementation between BBS-2 and IFT-A subunit IFT43 in *bbs-2(ok2053)* animals. IFT70/DYF-1::mScarlet was co-expressed as a ciliary IFT marker. *bbs-2(ok2053)* animals expressing IFT70/DYF-1::mScarlet alone are shown as controls. Asterisks and arrowheads are as in (D). Scale bar, 2.5 μm. **(F)** Representative ciliary images showing split-GFP complementation between BBS-7 and IFT-A subunit IFT43 in *bbs-7(n1606)* animals. IFT70/DYF-1::mScarlet was co-expressed as a ciliary IFT marker. *bbs-7(n1606)* animals expressing IFT70/DYF-1::mScarlet alone are shown as controls. Asterisks and arrowheads are as in (D). Scale bar, 2.5 μm. **(G)** Quantification of relative IFT70/DYF-1::mScarlet fluorescence intensity in the control and split-GFP rescue conditions shown in (D–F), calculated as the distal-to-middle segment fluorescence ratio. Each dot represents one cilium. N = 10 cilia per genotype. Data are mean ± s.d. n.s., not significant; ****P < 0.0001, two-tailed Student’s t-tests for the indicated comparisons. **(H)** Dye-filling rescue assay for the split-GFP-tagged constructs shown in (D–F). For each genotype, three independent plates were analyzed, with 100 animals scored per plate. Data are mean ± s.d. ****P < 0.0001, two-tailed Student’s t-tests for the indicated comparisons.

We initially attempted to test candidate BBSome – IFT proximities by FLIM-FRET using full-length fluorescently tagged protein pairs. However, although singly tagged reporters such as mScarlet::BBS-8 and GFP::IFT43 were detectable throughout cilia, co-expression of candidate BBSome – IFT pairs, including mScarlet::BBS-8 with IFT52::GFP and BBS-2::mScarlet with GFP::IFT43, altered the ciliary enrichment and segmental distribution of the tagged proteins (Fig. S4E,F). These observations suggested that bulky fluorescent proteins may perturb native BBSome – IFT arrangements, possibly because these candidate proximity sites are spatially constrained within IFT trains.

To overcome this limitation, we used split-GFP complementation as an *in vivo* proximity assay by fusing candidate partners to GFP1-10 and GFP11 fragments. Because split-GFP complementation can stabilize weak or transient contacts [41], we interpreted this assay as a topology and proximity readout rather than a direct measurement of binding affinity. We first benchmarked the specificity of the split-GFP assay using control pairs. As an internal positive control, IFT46/DYF-6 and IFT52/OSM-6 produced the expected split-GFP signal (Fig. S4G), consistent with their close association within the IFT-B complex [22] and with our FLIM-FRET results (Fig. 2M – P). As a structure-guided negative control, we tested the IFT-A subunit IFT144/DYF-2 and the IFT-B subunit IFT52/OSM-6, which are positioned in spatially separated regions of the anterograde IFT train architecture [22]. This pair did not produce detectable split-GFP signal under the same conditions (Fig. S4H), supporting the interpretation that split-GFP complementation in this setting reports spatially constrained proximity rather than nonspecific ciliary co-localization.

We then tested candidate BBSome – IFT proximities. For the IFT-B-associated site, structural modeling suggested that the IFT46 – IFT52 C-terminal module lies near the surface of the assembled BBSome. We therefore performed an initial split-GFP proximity test between BBS-8 and the IFT-B subunits IFT46/DYF-6 or IFT52/OSM-6. Split-GFP complementation was detected for both pairs in cilia, supporting close spatial proximity between the BBSome and the IFT46 – IFT52-containing IFT-B core (Fig. 5D). Because BBS-8 was the only BBSome subunit tested for this IFT-B-associated site, this result identifies BBSome proximity to the IFT46 – IFT52 module but does not define BBS-8 as the principal binding interface or exclude contributions from other BBSome subunits. For the IFT-A-associated candidate site, AlphaFold 3 models placed the N-terminal region of IFT43 within the BBSome, but the predicted local arrangement varied among models, suggesting conformational flexibility. We therefore tested IFT43 against multiple BBSome subunits. Split-GFP complementation was detected with BBS-2 and BBS-7 (Fig. 5E,F), whereas BBS-4, BBS-5, and BBS-9 did not produce detectable signal under the same conditions (Fig. S4I), consistent with a restricted BBSome proximity-accessible site near IFT43.

To determine whether the split-GFP-complemented BBSome – IFT pairs retained functional relevance *in vivo*, we examined IFT distribution and sensory cilium function in the corresponding mutant backgrounds. Expression of the split-GFP-tagged BBSome – IFT pairs reduced abnormal distal-segment accumulation of the IFT-B marker IFT70/DYF-1::mScarlet, as quantified by the distal-to-middle segment fluorescence ratio, and rescued dye-filling defects (Fig. 5D – H). These rescue data indicate that the split-GFP-tagged complexes remained compatible with ciliary IFT organization and sensory cilium function; they are consistent with a physiologically relevant proximity relationship.

Together, these split-GFP and rescue data indicate that the BBSome can be positioned in close proximity to both IFT-B-associated and IFT-A-associated sites in a functionally compatible state. Because split-GFP complementation can stabilize spatially proximal proteins and reports proximity rather than binding affinity or interface chemistry, these data do not by themselves define the molecular contact surface. Instead, these multivalent proximities motivate a working model in which the BBSome contributes to IFT-A – IFT-B coordination during anterograde transport.

### Conserved charged surfaces nominate candidate BBSome – IFT-B electrostatic complementarity

The multivalent BBSome – IFT proximity map raised the question of how the BBSome associates with the IFT complex. BBS-8::mScarlet was reduced in the residual ciliary compartment of *ift46(Q370*)* and *ift52(Q377*)* mutants, which carry C-terminal truncations of IFT46/DYF-6 and IFT52/OSM-6, respectively, and form shortened cilia lacking the distal segment (Fig. S5A). Because these truncations may broadly affect IFT-B assembly, stability, or cilium structure, this result does not specifically identify a BBSome docking site. By contrast, BBS-8::mScarlet remained detectable in cilia of *ift43(cas530)* mutants (Fig. S5B). These observations motivated examination of the IFT46 – IFT52 C-terminal module as a candidate contributor to BBSome recruitment.

We therefore focused on the IFT46 – IFT52 module identified by split-GFP complementation as a BBSome-proximal IFT-B site. AlphaFold 3 – based modeling, aligned to the available IFT-B structural framework [18][22], predicted that the C-terminal regions of IFT46/DYF-6 and IFT52/OSM-6 form a compact module positioned at the periphery of the IFT-B complex (Fig. 6A). Sequence inspection and surface electrostatic calculations revealed that this module contains an acidic surface patch generated by clustered glutamate and aspartate residues in the C-terminal regions of both IFT46 and IFT52 (Fig. 6B,C). Thus, the BBSome-proximal IFT46 –IFT52 module contains an acidic surface patch that could support charge-dependent interactions with the BBSome.

**Figure 6.**
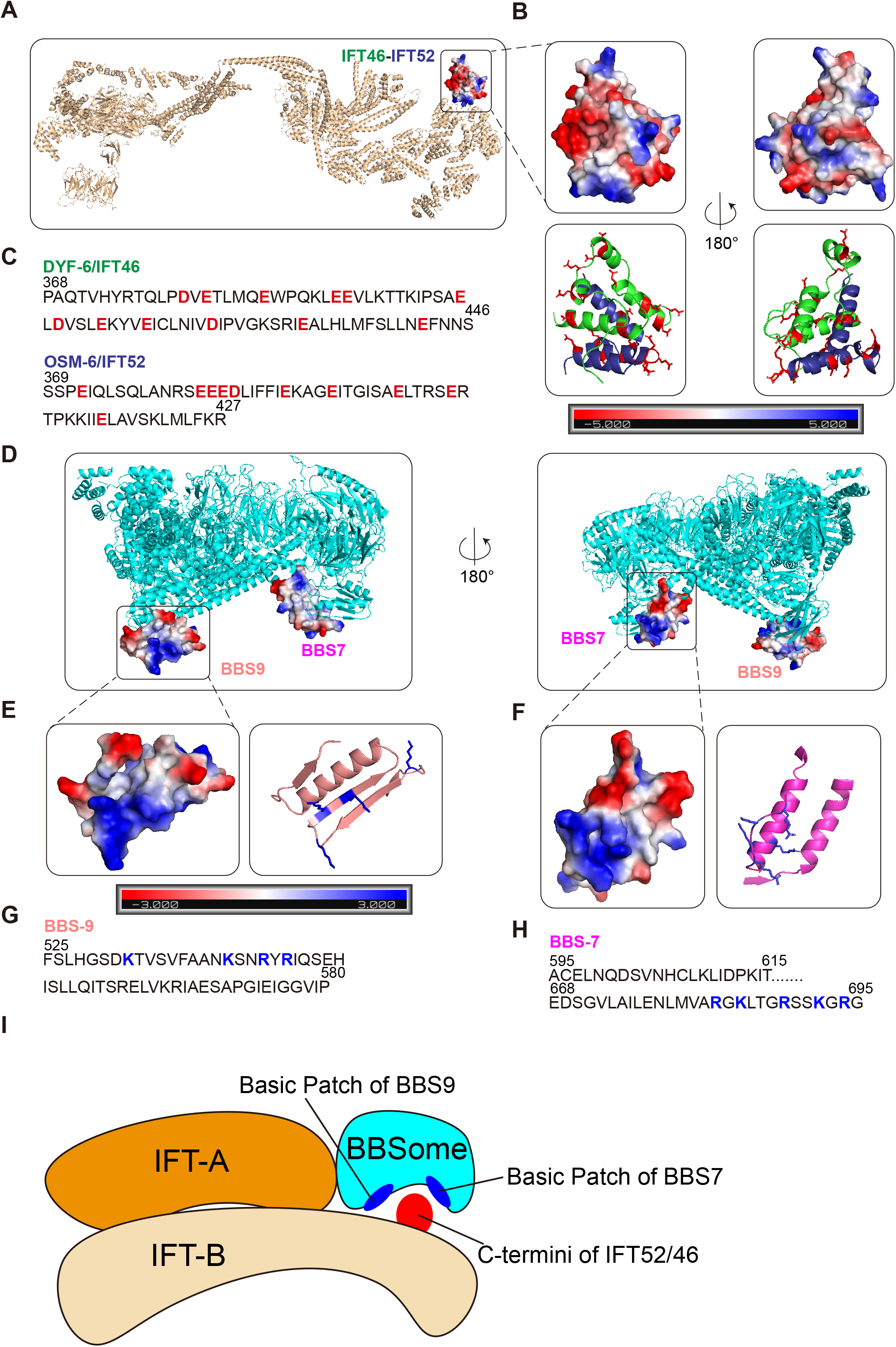
Electrostatic complementarity nominates a candidate BBSome–IFT-B interface. (A) Structural model showing the position of the C-terminal IFT46/DYF-6–IFT52/OSM-6 module within the IFT-B complex. The C-terminal regions of *C. elegans* IFT46/DYF-6 and IFT52/OSM-6 were modeled using AlphaFold 3 and aligned to the reported IFT-B structure (8BD7). The boxed region indicates the IFT46–IFT52 module analyzed for surface electrostatic potential. Surface electrostatic potential was calculated using the APBS plugin in PyMOL and displayed on a scale from −5 to +5 kBT/e. **(B)** Close-up views of the predicted IFT46–IFT52 C-terminal module. Upper panels show surface electrostatic potential, and lower panels show the corresponding structural model with acidic side chains highlighted in red. The right panels show the same module after 180° rotation. **(C)** Amino-acid sequences of the IFT46/DYF-6 and IFT52/OSM-6 C-terminal regions analyzed in (A and B). Acidic residues are highlighted in red. **(D)** Surface electrostatic potential of candidate BBSome regions containing BBS-9 and BBS-7. The *C. elegans* BBSome model was predicted using AlphaFold3 and aligned to the reported bovine BBSome structure (6VNW). Boxed regions indicate candidate electropositive surfaces. The right panel shows the same BBSome model after 180° rotation. Surface electrostatic potential was calculated using the APBS plugin in PyMOL and displayed on a scale from −3 to +3 kBT/e. **(E)** Close-up view of the candidate BBS-9 surface. Left, surface electrostatic potential; right, structural model with basic side chains highlighted in blue. **(F)** Close-up view of the candidate BBS-7 surface. Left, surface electrostatic potential; right, structural model with basic side chains highlighted in blue. **(G)** Amino-acid sequence of the BBS-9 region analyzed in (D and E). Basic residues are highlighted in blue. **(H)** Amino-acid sequence of the BBS-7 regions analyzed in (D and F). Basic residues are highlighted in blue. **(I)** Proposed working model positioning the BBSome relative to the anterograde IFT-A–IFT-B train architecture.

We next examined whether BBSome regions positioned near the IFT46 – IFT52 module contain complementary electropositive surfaces. In AlphaFold 3 – based composite models of the *C. elegans* BBSome with the IFT46 – IFT52 C-terminal module, three BBSome subunits—BBS-1, BBS-7, and BBS-9—were positioned near this acidic IFT-B module (Fig. S5C). Surface electrostatic analysis identified exposed basic patches on BBS-7 and BBS-9 within the assembled BBSome (Fig. 6D – H). These patches are enriched in lysine and arginine residues and are spatially compatible with the acidic surface of the IFT46 – IFT52 module in a proposed working placement of the BBSome relative to the anterograde IFT-A – IFT-B train architecture (Fig. 6I). A candidate electropositive BBS-1 surface was also positioned near the IFT46 – IFT52 C-terminal module in the composite model (Fig. S7A – C). Together, these structural analyses identify BBS-1, BBS-7, and BBS-9 as candidate BBSome surfaces that may engage the acidic IFT46 – IFT52 module.

To ask whether this electrostatic organization is evolutionarily conserved, we compared the corresponding IFT-B and BBSome regions across metazoan and algal species. The acidic character of the IFT46–IFT52 C-terminal module was retained across mammalian, *C. elegans*, and *C. reinhardtii* sequences and structural models (Fig. S6A–C). Conversely, the candidate BBSome regions positioned near the IFT46–IFT52 C-terminal module retained clusters of basic residues across species (Figs. S6D and S7C). Surface electrostatic calculations further showed that the electropositive features of the BBS-7/BBS-9 region are preserved in the experimentally determined bovine BBSome structure [39][40] and in predicted human and *C. reinhardtii* models (Fig. S6E,F). Together, these analyses nominate conserved electrostatic features consistent with candidate complementarity between the IFT46 –IFT52 module and BBS-1, BBS-7, or BBS-9, providing a testable structural framework for BBSome association with the IFT-B core.

### Charge reversal of candidate charged surfaces alters BBSome recruitment and ciliary function

The conserved acidic – basic surface complementarity between the IFT46 – IFT52 module and candidate basic surfaces on BBSome subunits suggested a potential electrostatic mechanism for BBSome engagement with IFT-B. To test whether these charged surfaces are functionally important *in vivo*, we generated transgenic charge-reversal variants targeting both candidate charged surfaces. Acidic residues within the C-terminal regions of IFT46/DYF-6 and IFT52/OSM-6, marked red in Fig. 6C, were converted to basic residues, whereas basic residues within candidate BBSome surfaces, marked blue in Fig. 6G,H and Fig. S7C, were converted to acidic residues.

We first established rescue conditions using wild-type constructs. In *ift46(Q370*)* animals, expression of wild-type IFT46/DYF-6::mScarlet restored ciliary localization of BBS-7::GFP. Similarly, in *ift52(Q377*)* animals, wild-type IFT52/OSM-6::GFP restored ciliary localization of BBS-9::mScarlet (Fig. 7A,B,E). These wild-type constructs also rescued dye-filling defects, indicating functional restoration of sensory cilia (Fig. 7G). By contrast, charge-reversal variants of IFT46 or IFT52 markedly reduced ciliary enrichment of BBS-7 or BBS-9, respectively, despite the presence of ciliary IFT markers (Fig. 7A,B,E). These variants also failed to efficiently rescue dye-filling, indicating that extensive charge reversal of the acidic C-terminal surfaces of IFT46 and IFT52 compromises BBSome enrichment and sensory cilium function under these rescue conditions (Fig. 7G).

**Figure 7.**
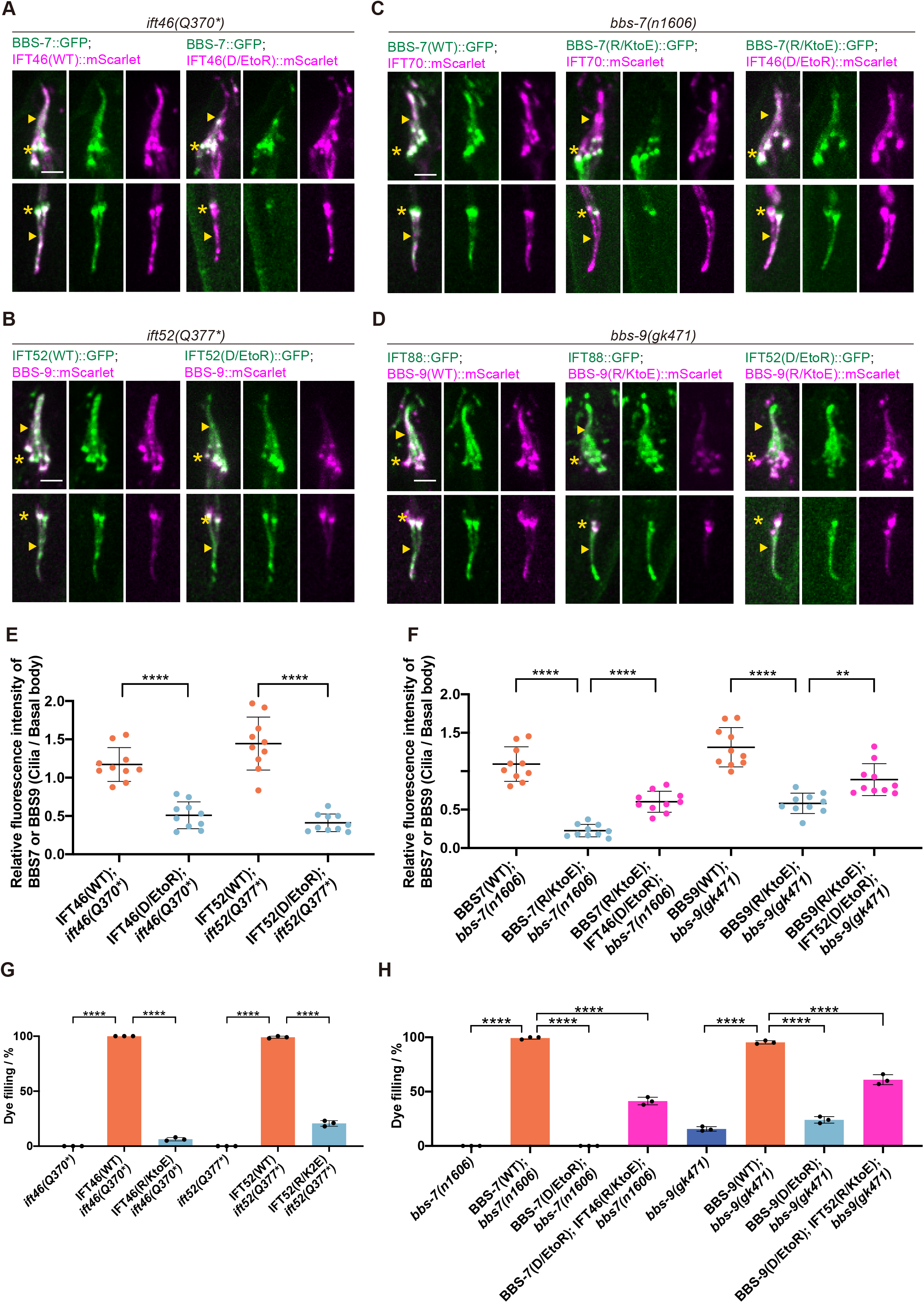
Charge-reversal and reciprocal charge-swap experiments test candidate charged surfaces involved in BBSome recruitment and ciliary function. (A) Representative ciliary images of *ift46(Q370*)* animals expressing BBS-7::GFP together with either wild-type IFT46::mScarlet or the IFT46(D/EtoR)::mScarlet variant. IFT46(D/EtoR) denotes a charge-reversal variant in which all the acidic residues highlighted in red in the IFT46 C-terminal region in Figure 6C were substituted with arginine. Asterisks mark the ciliary base/transition zone, and arrowheads mark the junction between the middle and distal segments. Scale bar, 2.5 μm. **(B)** Representative ciliary images of *ift52(Q377*)* animals expressing BBS-9::mScarlet together with either wild-type IFT52::GFP or the IFT52(D/EtoR)::GFP variant. IFT52(D/EtoR) denotes a charge-reversal variant in which all the acidic residues highlighted in red in the IFT52 C-terminal region in Figure 6C were substituted with arginine. Asterisks and arrowheads are as in (A). Scale bar, 2.5 μm. **(C)** Representative ciliary images of *bbs-7(n1606)* animals expressing wild-type BBS-7::GFP, the charge-reversal BBS-7(R/KtoE)::GFP variant, or BBS-7(R/KtoE)::GFP together with the reciprocal charge-reversal IFT46(D/EtoR)::mScarlet variant. BBS-7(R/KtoE) denotes a charge-reversal variant in which all the basic residues highlighted in blue in Figure 6H were substituted with glutamate. In the first two conditions, IFT70::mScarlet was co-expressed as a ciliary IFT marker. Asterisks and arrowheads are as in (A). Scale bar, 2.5 μm. **(D)** Representative ciliary images of *bbs-9(gk471)* animals expressing wild-type BBS-9::mScarlet, the charge-reversal BBS-9(R/KtoE)::mScarlet variant, or BBS-9(R/KtoE)::mScarlet together with the reciprocal charge-reversal IFT52(D/EtoR)::GFP variant. BBS-9(R/KtoE) denotes a charge-reversal variant in which all basic residues highlighted in blue in Figure 6G were substituted with glutamate. In the first two conditions, IFT88::GFP was co-expressed as a ciliary IFT marker. Asterisks and arrowheads are as in (A). Scale bar, 2.5 μm. **(E)** Quantification of BBS-7::GFP or BBS-9::mScarlet ciliary enrichment in the wild-type and charge-reversal IFT46 or IFT52 rescue conditions shown in (A) and (B). Ciliary enrichment was calculated as the fluorescence intensity in the ciliary shaft divided by the fluorescence intensity at the basal body. Ten phasmid cilia were analyzed per condition. Data are mean ± s.d. ****P < 0.0001, one-way ANOVA followed by Tukey’s multiple-comparisons test. **(F)** Quantification of BBS-7::GFP or BBS-9::mScarlet ciliary enrichment in the wild-type, charge-reversal, and reciprocal charge-swap conditions shown in (C) and (D). Ciliary enrichment was calculated as in (E). Ten phasmid cilia were analyzed per condition. Data are mean ± s.d. **P < 0.01; ****P < 0.0001, one-way ANOVA followed by Tukey’s multiple-comparisons test. **(G)** DiI dye-filling rescue assay for wild-type and charge-reversal IFT46 and IFT52 constructs in the corresponding IFT mutant backgrounds. For each genotype, three independent plates were analyzed, with 100 animals scored per plate. Data are mean ± s.d. ***P < 0.001; ****P < 0.0001, one-way ANOVA followed by Tukey’s multiple-comparisons test. **(H)** DiI dye-filling rescue assay for the wild-type, charge-reversal, and reciprocal charge-swap BBS-7 and BBS-9 conditions in the corresponding BBSome mutant backgrounds. For each genotype, three independent plates were analyzed, with 100 animals scored per plate. Data are mean ± s.d. *P < 0.05; ****P < 0.0001, one-way ANOVA followed by Tukey’s multiple-comparisons test.

We next tested the reciprocal charged surfaces on the BBSome. Because structural modeling identified candidate electropositive surfaces on BBS-1, BBS-7, and BBS-9, we first tested whether charge reversal of the BBS-1 basic patch affected BBSome ciliary entry. In *bbs-1(ok1111)* animals, wild-type BBS-1::GFP entered cilia and restored IFT70/DYF-1::mScarlet distribution. The BBS-1(R/KtoE)::GFP variant also entered cilia and did not reduce BBS-1 ciliary enrichment; instead, it showed increased cilia-to-basal body fluorescence relative to wild-type BBS-1 (Fig. S7D – E). Thus, charge reversal of this candidate BBS-1 basic surface does not impair BBS-1 entry into cilia. This result suggests that the BBS-1 basic patch is not the limiting determinant for BBSome ciliary recruitment, although it may influence BBSome residence, accumulation, or recycling.

We therefore focused on the candidate complementary charged surfaces of the acidic IFT46 – IFT52 C-terminal module and basic regions of BBS-7 and BBS-9. Wild-type BBS-7::GFP and BBS-9::mScarlet restored ciliary localization and rescued dye-filling defects in the corresponding *bbs-7* and *bbs-9* mutant backgrounds (Fig. 7C, D, F and H). In contrast, BBS-7(R/KtoE)::GFP and BBS-9(R/KtoE)::mScarlet showed substantially reduced ciliary enrichment relative to their wild-type counterparts and failed to restore normal dye-filling (Fig. 7C, D, F and H). Thus, charge reversal of either the acidic IFT-B surface or the basic BBS-7/BBS-9 surfaces compromises BBSome localization and ciliary function.

To test whether electrostatic compatibility contributes to these phenotypes, we performed reciprocal charge-swap experiments. Co-expression of BBS-7(R/KtoE)::GFP with IFT46(D/EtoR)::mScarlet, or BBS-9(R/KtoE)::mScarlet with IFT52(D/EtoR)::GFP, partially restored BBSome ciliary enrichment and dye-filling compared with the BBS-7 or BBS-9 charge-reversal variants alone (Fig. 7C, D, F and H). This partial suppression is consistent with a contribution of electrostatic compatibility to BBSome recruitment. However, the unequal sets of substituted residues, multicopy transgene expression, and possible differences in array composition preclude assignment of a specific pairwise interaction. These experiments therefore support a functional role for the candidate charged surfaces without establishing a direct BBSome – IFT-B interface.

## Discussion

How IFT-A and IFT-B remain coordinated within dynamic ciliary transport trains is a fundamental question in cilia biology. Here, we combine endogenous fluorescent knock-in labeling, *in vivo* FLIM-FRET, structural modeling, and targeted charge-reversal mutagenesis to test a working model for BBSome-associated IFT-A –IFT-B coordination. Our data are consistent with multivalent BBSome proximity to IFT-A- and IFT-B-associated sites, including candidate complementary charged surfaces on the IFT46 – IFT52 module and BBS7/BBS9. This framework links subunit proximity, transport dynamics, and ciliary function within living cilia.

Endogenous FLIM-FRET provides an *in vivo* proximity readout, and the agreement of selected IFT-B pairs with available structures supports assay validity under the conditions tested. However, whole-cilium mono-exponential fits yield an apparent, amplitude-weighted mean lifetime from mixed FRET-active and non-FRET donor populations. They do not resolve FRET efficiency or the FRET-active fraction, and lifetime shifts can depend on acceptor abundance and spatial co-occupancy. Thus, reduced lifetime shifts are consistent with altered proximity or co-occupancy, but cannot by themselves define loss of a specific local interface or its directionality.

Our findings refine and extend previous work implicating the BBSome in IFT organization. The BBSome has been best characterized as a ciliary trafficking complex that mediates the transport and recycling of membrane-associated cargoes [42][43][44]. Earlier studies in *C. elegans* showed that BBSome disruption separates IFT-A and IFT-B transport behavior, suggesting a role in maintaining IFT-train integrity [29][33]. However, those studies relied largely on transgenic fluorescent reporters and did not directly assess molecular proximity between IFT subcomplexes. By revisiting this question with endogenous knock-in strains and FLIM-FRET, our study shows that BBSome loss is associated with reduced whole-cilium IFT-A – IFT-B FLIM-FRET signals. Mammalian IMCD3 cell experiments further support this conclusion: although IFT144 and IFT88 both enter BBS7-knockout cilia, their transport velocities diverge and their FLIM-FRET proximity is reduced. These results are consistent with a similar contribution of the BBSome to IFT-A – IFT-B coordination across the two systems, while same-cell dual-reporter transport measurements would strengthen this conclusion.

Mechanistically, our data are consistent with the BBSome contributing to IFT coordination through distributed proximity relationships rather than a single rigid binding site. Split-GFP proximity assays place BBSome components near both IFT-A- and IFT-B-associated components, including IFT43 and the IFT46 – IFT52 module. Structural modeling further nominates candidate electrostatic complementarity between acidic C-terminal surfaces of IFT46 and IFT52 and basic surfaces on BBS7 and BBS9. Charge-reversal mutations targeting these candidate surfaces reduce BBSome ciliary enrichment and fail to restore sensory cilium function, supporting the functional importance of these charged surfaces *in vivo*. Such distributed electrostatic contributions may be well suited to the IFT system, which must maintain train cohesion while allowing continuous remodeling, cargo exchange and directional switching [7].

Our working model for BBSome localization within IFT trains is broadly consistent with previous studies of BBSome – IFT interplay [45]. The C-terminal tail of IFT38 has been shown to interact directly with the BBSome [39][46]. This tail is unresolved in current IFT-train structures; therefore, its spatial relationship to the IFT46 – IFT52 module remains uncertain. Nevertheless, the IFT46 – IFT52 C-terminal module occupies a peripheral position within IFT-B1 [22], consistent with accessibility at the BBSome-proximal site proposed here. IFT22/RABL5 recruits the BBSome to the basal body through ARL6/BBS3 [47], whereas IFT25 – IFT27 participates in BBSome-dependent ciliary trafficking [42][37][48][49]. IFT22 and IFT25 – IFT27 associate with the IFT74 – IFT81 module within IFT-B [18][50], and this module is also positioned near the BBSome in our working model. Together, these observations support a BBSome location adjacent to multiple IFT-B modules but do not establish a direct binding interface.

This model also provides a broader conceptual view of BBSome function. Rather than acting solely as a cargo adaptor or coat-like sorting complex, the BBSome may also contribute to the organization of the IFT machinery. Through proximity to multiple IFT-associated sites, the BBSome may help stabilize the relative positioning of IFT-A and IFT-B during transport. Such a coordinating role could explain why BBSome mutants show defects not only in cargo trafficking but also in IFT train behavior, subcomplex distribution and ciliary organization [32][33]. In this sense, cargo trafficking and train cohesion may be mechanistically linked: the same multivalent surfaces that position the BBSome within IFT trains may also create a platform for regulated cargo loading, retention and release.

Several aspects of the model remain to be resolved. FLIM-FRET and split-GFP provide powerful readouts of molecular proximity in intact cilia, but they do not by themselves define atomic binding interfaces or binding affinities. Similarly, AlphaFold-based modeling and electrostatic surface analysis nominate plausible interaction surfaces, but high-resolution structures of BBSome-associated IFT trains will be required to define the interface at atomic resolution. Charge-reversal mutations provide functional support for the importance of the candidate charged surfaces, but these mutations could also affect local folding, complex assembly or protein stability. Although reciprocal charge-swap experiments yielded partial suppression, the substitutions involve unequal residue sets, and the multicopy rescue arrays may differ in expression or composition. These results do not establish direct residue pairing; future biochemical reconstitution and in situ structural analysis will be important to distinguish electrostatic pairing from broader effects on BBSome–IFT assembly.

Together, our findings support a working model in which multivalent BBSome – IFT proximities contribute to IFT-A – IFT-B coordination *in vivo*. This work extends the view of the BBSome beyond cargo trafficking and identifies candidate relationships that can be tested by biochemical and structural approaches.

## Acknowledgments

We thank the Imaging Core Facility, Technology Center for Protein Science, Tsinghua University, for assistance with FV1200 confocal microscopy. We also thank Bingyu Liu for guidance with image analysis. This work was supported by the National Natural Science Foundation of China grants 31991190, 32430026, 32270721, 32470730, 32270773, and 92254306; the National Key Research and Development Program of China grants 2024YFA1307301, 2022YFA1302700, and 2019YFA0508401; the Pillars of the Nation Funding for Life Sciences, Tsinghua University, and TsienTang Life Science Development Fund at Tsinghua University.

## Author contributions

G. Ou and Z. Shen conceived the project. Z. Shen performed the experiments and collected and analyzed the data under the supervision of G. Ou. G. Chen and M. Li contributed to structural modeling of the BBSome and anterograde IFT train. G. Ou, Z. Shen, and W. Li wrote the manuscript.

## Data and materials availability

Data supporting the findings are provided in the main text and supplementary materials. Raw imaging and analysis files are available from the corresponding author upon reasonable request.

## Supplementary Materials

### Materials and Methods

Supplementary Figures S1–S7

### Methods Molecular Biology

*C. elegans* cDNAs for *bbs-7*, *bbs-9*, *dyf-6* (IFT46), *osm-6* (IFT52), and the other indicated genes were cloned into pDONR plasmids with N- or C-terminal tags using the In-Fusion HD Cloning Kit (TaKaRa) according to the manufacturer’s instructions. Mouse *Ift144* and *Ift88* cDNAs were cloned into pcDNA3.0 for transfection or into pLV for lentiviral production using In-Fusion. Mutant constructs were generated by PCR. All plasmid inserts were verified by Sanger sequencing.

### *C. elegans* culture and genetics

*C. elegans* were cultured at 20 °C on standard NGM (nematode growth medium) plates seeded with OP50. Knock-in strains of fluorescence-tagged IFT subunits were generated by Sunybiotech through CRISPR-Cas9. Double-tagged strains of IFT subunits were generated by genetic crossing between two knock-in strains and confirmed by confocal microscopy and PCR. All single- and double-tagged knock-in strains used for FLIM were homozygous at the tagged loci, as verified by PCR.

For transgenic strains, constructs of WT or mutant genes were co-injected with a plasmid (pRF4) encoding *rol-6*[*su1006*] as the marker into the germ line of young adult hermaphrodites. Roller F1s were singled and the F2s with transgenes were identified as transgenic lines. At least two independent lines were maintained for each strain; lines showed comparable phenotypes and were pooled for analysis.

### Cell culture

IMCD3 cells were cultured in DMEM/F12 supplemented with 10% FBS and 1% penicillin–streptomycin at 37 °C and 5% CO_2_. IFT144 and IFT88 plasmids were transfected using Lipofectamine 3000 (Thermo Fisher Scientific) according to the manufacturer’s instructions.

### Lentivirus production and infection

2μg pLV plasmids containing BBS7 sgRNA plus Cas9 or fluorescence-tagged IFT subunits were first transfected into HEK293T cells using Lipofectamine 3000, together with 1.5μg packaging plasmid psPAX2 and 1μg envelope plasmid PMD2.G.

After 48 h, supernatant containing mature lentivirus was collected and filtered with 0.22 μm filter.

IMCD3 cells were infected with lentivirus and cultured for another 48 h. The cells were first selected for puromycin or neomycin resistance by culturing under 10 μg/mL puromycin or 200 μg/mL neomycin for 48 hours. Then, single-cell colonies were selected. For BBS7 KO cells, each colony was first analyzed by Sanger sequencing and further tested by SMO immunostaining. For cells stably expressing fluorescence-tagged IFT subunits, colonies were verified under confocal microscopy.

### Immunostaining

IMCD3 cells were cultured in glass-bottom plates and serum-starved for >24 h to induce ciliogenesis. Cells were washed three times with PBS and fixed with 4% paraformaldehyde. After three washes with PBS, cells were blocked with 5% BSA in PBSTX (PBS containing 0.1% Triton X-100) for 1 h at room temperature and then incubated overnight at 4 °C with rabbit anti-SMO (AF8010; Beyotime) and mouse anti-ARL13B (66739-1-Ig; Proteintech). After five washes with PBS, cells were incubated for 1 h at room temperature with DyLight 488 goat anti-rabbit IgG (A23220; Abbkine) and DyLight 549 goat anti-mouse IgG (A23310; Abbkine), followed by five washes with PBS before imaging.

### Live imaging

Live imaging of *C. elegans* was performed as described previously [51]. Young adult worms were anesthetized with 0.1 mmol/L levamisole in M9 buffer and immobilized on 3% agarose pads. Imaging was performed on an Axio Observer Z1 microscope (Carl Zeiss) equipped with a 100×, 1.49-NA objective and an iXon+ DU-897D-C00#BV-500 electron-multiplying charge-coupled device camera (Andor). A Sapphire CW CDRH USB laser system provided 488- and 561-nm illumination through a CSU-X1 spinning-disk confocal scan head (Yokogawa). Images were acquired with μManager [52] at 200-ms exposure and analyzed in Fiji [53]. For each cilium, a 40-s movie was acquired to generate kymographs and measure IFT velocities. The middle–distal boundary was identified in the fluorescence image at the junction between the wider middle segment and the narrower distal segment; kymographs were generated separately for the two segments using this boundary. IMCD3 cells were cultured in glass-bottom plates and serum-starved for 24 h to induce ciliogenesis. Imaging and analysis were then performed as described above for C. elegans.

#### FLIM

For fluorescence lifetime imaging, young adult worms were anesthetized and immobilized as described for live imaging, and IMCD3 cells were serum-starved as described above. FLIM was performed on an inverted FV1200 microscope (Olympus) equipped with a PicoHarp 300 controller (PicoQuant). Excitation at 488 nm was provided by a picosecond pulsed diode laser at a repetition rate of 40 MHz through a 100×, 1.49 NA objective. Emission was filtered with a 510/55-nm bandpass filter and detected with a single-photon avalanche diode detector (Micro Photon Devices). Whole-cilium regions of interest were manually selected from fluorescence intensity images, and at least 50,000 photons were acquired per region of interest. The fluorescence decay curve for each cilium was fitted in SymPhoTime 64 (PicoQuant) with a mono-exponential decay model. For mixed FRET-active and non-FRET donor populations, the fitted value represents an apparent, amplitude-weighted mean lifetime rather than a species-specific lifetime.

### Dye-filling assay

Young adult worms were collected from NGM plates in M9 buffer. DiI (1,1′-dioctadecyl-3,3,3′,3′-tetramethylindocarbocyanine perchlorate; Sigma-Aldrich) was added to a final concentration of 20 μg/ml. After incubation in the dark for 45 min at room temperature, worms were transferred to NGM plates seeded with OP50. Dye-filling was scored 2 h later using a fluorescence stereomicroscope.

### Quantifications and statistical analysis

ImageJ was used for image measurements and quantification. For IFT velocity, individual transport events were identified in kymographs from at least 10 independent cilia per group. Event-level velocity distributions and Gaussian fits were used for visualization; inferential comparisons were performed using the mean velocity for each cilium as the independent replicate. FLIM analyses used one apparent mean lifetime per cilium; unless otherwise noted, 10 cilia from 10 animals or cells were analyzed per condition. For dye-filling assays, each independently scored plate was treated as one replicate. Statistical analyses were performed in GraphPad Prism, and the tests are specified in the figure legends.

### AlphaFold 3 prediction

IFT and BBSome structures were predicted using the AlphaFold Server (AlphaFold 3) [54]. The top-ranked models were analyzed in PyMOL. Surface electrostatic potentials were calculated with the APBS plugin at pH 7.0, protein dielectric 2.0, solvent dielectric 78.0, and ionic strength 150 mM; protonation states were assigned using PDB2PQR [55]. Models were used for hypothesis generation rather than as evidence of direct binding.

### Materials Bacterial strains

*Escherichia coli* (OP50), CGC, OP50

### Chemicals

Levamisole, Sigma-Aldrich, Cat# L9756 DMEM/F12, Gibco, Cat# 11320033 FBS, Gibco, Cat# A5256701

### Critical commercial assays

In-Fusion HD Cloning Kit, TaKaRa, Cat# 639650 Lipofectamine 3000, Thermo Fisher, Cat# L3000015

### Antibodies

Rabbit anti-SMO, Beyotime, Cat# AF8010

Mouse anti-ARL13B, Proteintech, Cat# 66739-1-Ig Dylight 488 Goat Anti-Rabbit IgG, Abbkine, Cat# A23220 Dylight 549 Goat Anti-Mouse IgG, Abbkine, Cat# A23310

**Experimental models: Cell lines** Mouse: mIMCD3, ATCC, CRL-2123 Human: HEK-293T, ATCC, CRL-3216

### Experimental models: Organisms/strains

*C. elegans*:

PHX7530: *syb7530*[*osm-6::GFP*]

PHX7487: *syb7487*[*dyf-6::mScarlet*] PHX7160: *syb7160*[*dyf-3::mScarlet*] PHX7521: *syb7521*[*bbs-9::GFP*] PHX7122: *syb7122*[*bbs-7::GFP*]

PHX7723: *syb7723*[*ift-139::mScarlet*] PHX7439: *syb7439*[*osm-1::GFP*] PHX7466: *syb7466*[*osm-5::GFP*]

PHX7481: *syb7481*[*dyf-2::mScarlet*] PHX7648: *syb7648*[*ifta-2::GFP*] PHX7227: *syb7227*[*che-13::GFP*]

PHX7571: *syb7571*[*daf-10::mScarlet*] PHX7493: *syb7493*[*ift-43::GFP*]

PHX7548: *syb7548*[*ift-81::mScarlet*] PHX7611: *syb7611*[*dyf-13::GFP*]

PHX7478: *syb7478*[*dyf-1::mScarlet*] PHX7496: *syb7496*[*ifta-1::GFP*] GOU2360: *cas497*[*che-11::GFP*] GOU2362: *cas499*[*ift-74::GFP*] GOU4012: *cas1014*[*dyf-11::3xGFP*] GOU4010: *cas1013*[*che-2::3xGFP*]

GOU4772: *cas1086*[*dyf-11::mScarlet*]

GOU6001: *syb7160[dyf-3::mScarlet]*; *syb7227[che-13::GFP]* GOU6002: *cas1086[dyf-11::mScarlet]*; *cas497[che-11::GFP]* GOU6003: *syb7160[dyf-3::mScarlet]*; *cas499[ift-74::GFP]* GOU6005: *syb7160[dyf-3::mScarlet]*; *cas1014[dyf-11::3xGFP]* GOU6006: *syb7160[dyf-3::mScarlet]*; *cas1013[che-2::3xGFP]* GOU6007: *syb7160[dyf-3::mScarlet]*; *cas497[che-11::GFP]* GOU6008: *syb7160[dyf-3::mScarlet]*; *syb7439[osm-1::GFP]* GOU6009: *cas1086[dyf-11::mScarlet]*; *cas499[ift-74::GFP]* GOU6011: *cas1086[dyf-11::mScarlet]*; *cas1013[che-2::3xGFP]* GOU6012: *cas1086[dyf-11::mScarlet]*; *syb7439[osm-1::GFP]* GOU6013: *syb7160[dyf-3::mScarlet]*; *syb7493[ift-43::GFP]* GOU6014: *cas1086[dyf-11::mScarlet]*; *syb7227[che-13::GFP]* GOU6015: *syb7160[dyf-3::mScarlet]*; *syb7648[ifta-2::GFP]*

GOU6016: *cas1086[dyf-11::mScarlet]*; *syb7493[ift-43::GFP]* GOU6017: *cas1086[dyf-11::mScarlet]*; *syb7466[osm-5::GFP]* GOU6018: *syb7487[dyf-6::mScarlet]*; *syb7530[osm-6::GFP]* GOU6019: *syb7478[dyf-1::mScarlet]*; *syb7530[osm-6::GFP]* GOU6020: *syb7487[dyf-6::mScarlet]*; *syb7466[osm-5::GFP]* GOU6021: *syb7478[dyf-1::mScarlet]*; *syb7466[osm-5::GFP]* GOU6022: *syb7487[dyf-6::mScarlet]*; *syb7611[dyf-13::GFP]* GOU6023: *syb7478[dyf-1::mScarlet]*; *syb7611[dyf-13::GFP]* GOU6024: *syb7548[ift-81::mScarlet]*; *cas499[ift-74::GFP]* GOU6025: *syb7481[dyf-2::mScarlet]*; *syb7439[osm-1::GFP]* GOU6026: *syb7481[dyf-2::mScarlet]*; *syb7466[osm-5::GFP]*

GOU6040: *syb7481[dyf-2::mScarlet]*; *syb7439[osm-1::GFP]*; *bbs-7(n1606)* GOU6041: *syb7481[dyf-2::mScarlet]*; *syb7466[osm-5::GFP]*; *bbs-7(n1606)* GOU6042: *syb7481[dyf-2::mScarlet]*; *syb7439[osm-1::GFP]*; *bbs-1(ok1111)* GOU6043: *syb7481[dyf-2::mScarlet]*; *syb7439[osm-1::GFP]*; *bbs-8(nx77)* GOU6044: *syb7481[dyf-2::mScarlet]*; *syb7439[osm-1::GFP]*; *bbs-2(ok2053)* GOU6045: *syb7481[dyf-2::mScarlet]*; *syb7439[osm-1::GFP]*; *bbs-9(gk471)* GOU6046: *syb7478[dyf-1::mScarlet]*; *syb7521[bbs-9::GFP]*; *bbs-7(n1606)* MT3645: *bbs-7(n1606)*

GJ3457: *bbs-8(nx77)* VC5169: *bbs-2(ok2053)* VC1062: *bbs-9(gk471)* VC837: *bbs-1(ok1111)*

GOU6047: *cas1212[3xGFP::che-3]*; *dyf-6(Q370*)*

GOU6048: *cas1212[3xGFP::che-3]*; *osm-6(Q377*)*

GOU6049: *syb7481[dyf-2::mScarlet]*; *syb7122[bbs-7::GFP]*

GOU6051: *bbs-8(nx77)*; *casEx8011* [*Pdyf-1::osm-6::GFP1-10;*

*Pdyf-1::GFP11::bbs-8; Pdyf-1::dyf-1::mScarlet; pRF4(+)*]

GOU6052: *bbs-8(nx77)*; *casEx8012* [*Pdyf-1::dyf-6::GFP1-10; Pdyf-1::GFP11::bbs-8;*

*Pdyf-1::dyf-1::mScarlet; pRF4(+)*]

GOU6053: *bbs-2(ok2053)*; *casEx8013* [*Pdyf-1::bbs-2::GFP1-10;*

*Pdyf-1::GFP11::ift-43; Pdyf-1::dyf-1::mScarlet; pRF4(+)*]

GOU6054: *bbs-7(n1606)*; *casEx8014* [*Pdyf-1::bbs-7::GFP1-10;*

*Pdyf-1::GFP11::ift-43; Pdyf-1::dyf-1::mScarlet; pRF4(+)*] GOU6055: *casEx8015* [*Pdyf-1::mScarlet::bbs-8; pRF4(+)*]

GOU6056: *casEx8016* [*Pdyf-1::mScarlet::bbs-8; Pdyf-1::osm-6::GFP; pRF4(+)*] GOU6057: *casEx8017* [*Pdyf-1::GFP::ift-43; pRF4(+)*]

GOU6058: *casEx8018* [*Pdyf-1::GFP::ift-43; Pdyf-1::bbs-2::mScarlet; pRF4(+)*] GOU6059: *casEx8019* [*Pdyf-1::dyf-6::GFP1-10; Pdyf-1::osm-6::GFP11;*

*Pdyf-1::dyf-1::mScarlet; pRF4(+)*]

GOU6060: *casEx8020* [*Pdyf-1::dyf-2::GFP1-10; Pdyf-1::osm-6::GFP11;*

*Pdyf-1::dyf-1::mScarlet; pRF4(+)*]

GOU6062: *cas1212[3xGFP::che-3]*; *osm-6(Q377*)*; *casEx8022*

[*Pdyf-1::bbs-8::mScarlet; pRF4(+)*]

GOU6063: *cas1212[3xGFP::che-3]*; *dyf-6(Q370*)*; *casEx8023*

[*Pdyf-1::bbs-8::mScarlet; pRF4(+)*]

GOU6064: *dyf-6(Q370*)*; *casEx8024* [*Pdyf-1::dyf-6::mScarlet; Pdyf-1::bbs-7::GFP; pRF4(+)*]

GOU6065: *osm-6(Q377*)*; *casEx8025* [*Pdyf-1::osm-6::GFP;*

*Pdyf-1::bbs-9::mScarlet; pRF4(+)*]

GOU6066: *bbs-7(n1606)*; *casEx8026* [*Pdyf-1::dyf-1::mScarlet; Pdyf-1::bbs-7::GFP; pRF4(+)*]

GOU6067: *bbs-9(gk471)*; *casEx8027* [*Pdyf-1::osm-5::GFP; Pdyf-1::bbs-9::mScarlet; pRF4(+)*]

GOU6068: *dyf-6(Q370*)*; *casEx8028* [*Pdyf-1::dyf-6(D380R, E382R, E387R, E393R, E394R, E405R, D407R, E411R, E415R, D422R, E431R, E442R)::mScarlet; Pdyf-1::bbs-7::GFP; pRF4(+)*]

GOU6069: *osm-6(Q377*)*; *casEx8029* [*Pdyf-1::osm-6(E372R, E383R, E384R, E385R, D386R, E392R, E396R, E403R, E408R, E416R)::GFP; Pdyf-1::bbs-9::mScarlet; pRF4(+)*]

GOU6070: *bbs-7(n1606)*; *casEx8030* [*Pdyf-1::dyf-1::mScarlet; Pdyf-1::bbs-7(R683E, K685E, R689E, K692E, R694E)::GFP; pRF4(+)*]

GOU6071: *bbs-9(gk471)*; *casEx8031* [*Pdyf-1::osm-5::GFP; Pdyf-1::bbs-9(K532E,*

*K541E, R544E, R546E)::mScarlet; pRF4(+)*]

GOU6072: *bbs-7(n1606)*; *casEx8032* [*Pdyf-1::dyf-6(D380R, E382R, E387R, E393R, E394R, E405R, D407R, E411R, E415R, D422R, E431R, E442R)::mScarlet; Pdyf-1::bbs-7(R683E, K685E, R689E, K692E, R694E)::GFP; pRF4(+)*]

GOU6073: *bbs-9(gk471)*; *casEx8033* [*Pdyf-1::osm-6(E372R, E383R, E384R, E385R, D386R, E392R, E396R, E403R, E408R, E416R)::GFP; Pdyf-1::bbs-9(K532E, K541E, R544E, R546E)::mScarlet; pRF4(+)*]

GOU6074: *casEx8034* [*Pdyf-1::dyf-1::mScarlet; pRF4(+)*]

GOU6075: *bbs-8(nx77)*; *casEx8035* [*Pdyf-1::dyf-1::mScarlet; pRF4(+)*] GOU6076: *bbs-2(ok2053)*; *casEx8036* [*Pdyf-1::dyf-1::mScarlet; pRF4(+)*] GOU6077: *bbs-7(n1606)*; *casEx8037* [*Pdyf-1::dyf-1::mScarlet; pRF4(+)*]

GOU6078: *casEx8038* [*Pdyf-1::bbs-4::GFP1-10; Pdyf-1::GFP11::ift-43;*

*Pdyf-1::dyf-1::mScarlet; pRF4(+)*]

GOU6079: *casEx8039* [*Pdyf-1::bbs-5::GFP1-10; Pdyf-1::GFP11::ift-43;*

*Pdyf-1::dyf-1::mScarlet; pRF4(+)*]

GOU6080: *casEx8040* [*Pdyf-1::bbs-9::GFP1-10; Pdyf-1::GFP11::ift-43;*

*Pdyf-1::dyf-1::mScarlet; pRF4(+)*]

GOU6081: *bbs-1(ok1111)*; *casEx8041* [*Pdyf-1::dyf-1::mScarlet; Pdyf-1::bbs-1::GFP; pRF4(+)*]

GOU6082: *bbs-1(ok1111)*; *casEx8042* [*Pdyf-1::dyf-1::mScarlet; Pdyf-1::bbs-1(K127E, K132E, K133E, K144E, R145E, K154E, R164E, K167E, K180E)::GFP; pRF4(+)*]

GOU6083: *ift-43(cas530)*; *casEx8043* [*Pdyf-1::osm-5::GFP; Pdyf-1::bbs-8::mScarlet; pRF4(+)*]

### Plasmids

pDONR-*Pdyf-1::dyf-1::mScarlet*

pDONR-*Pdyf-1::dyf-6::GFP1-10* pDONR-*Pdyf-1::osm-6::GFP1-10* pDONR-*Pdyf-1::osm-6::GFP11*

pDONR-*Pdyf-1::mScarlet::bbs-8* pDONR-*Pdyf-1::bbs-8::mScarlet* pDONR-*Pdyf-1::GFP11::bbs-8*

pDONR-*Pdyf-1::bbs-2::mScarlet* pDONR-*Pdyf-1::bbs-2::GFP1-10* pDONR-*Pdyf-1::bbs-7::GFP1-10* pDONR-*Pdyf-1::bbs-4::GFP1-10* pDONR-*Pdyf-1::bbs-5::GFP1-10* pDONR-*Pdyf-1::bbs-9::GFP1-10* pDONR-*Pdyf-1::GFP::ift-43* pDONR-*Pdyf-1::GFP11::ift-43* pDONR-*Pdyf-1::dyf-2::GFP1-10* pDONR-*Pdyf-1::osm-5::GFP* pDONR-*Pdyf-1::osm-5::GFP11* pDONR-*Pdyf-1::bbs-7::GFP*

pDONR-*Pdyf-1::bbs-9::mScarlet* pDONR-*Pdyf-1::bbs-1::GFP* pDONR-*Pdyf-1::osm-6::GFP*

pDONR-*Pdyf-1::dyf-6::mScarlet*

pDONR-*Pdyf-1::bbs-7(R683E, K685E, R689E, K692E, R694E)::GFP*

pDONR-*Pdyf-1::bbs-9(K532E, K541E, R544E, R546E)::mScarlet*

pDONR-*Pdyf-1::osm-6(E372R, E383R, E384R, E385R, D386R, E392R, E396R,*

*E403R, E408R, E416R)::GFP*

pDONR-*Pdyf-1::dyf-6(D380R, E382R, E387R, E393R, E394R, E405R, D407R,*

*E411R, E415R, D422R, E431R, E442R)::mScarlet*

pDONR-*Pdyf-1::bbs-1(K127E, K132E,K133E, K144E, R145E, K154E, R164E, K167E, K180E)::GFP*

pLV-CMV::IFT144::EGFP

pLV-CMV::IFT88::mCherry pCDNA3.0-CMV::IFT144::3xNeoGreen pCDNA3.0-CMV::IFT88::3xNeoGreen

pLentiCRISPR V2-mouse-BBS7-sg(CGTGTCGAGCCCGCGTTGCT)

### Software

AlphaFold Server (AlphaFold 3), https://alphafoldserver.com/ Fiji https://fiji.sc/

GraphPad Prism 8 https://www.graphpad.com/ PyMOL https://pymol.org

---

**Figure S1.**
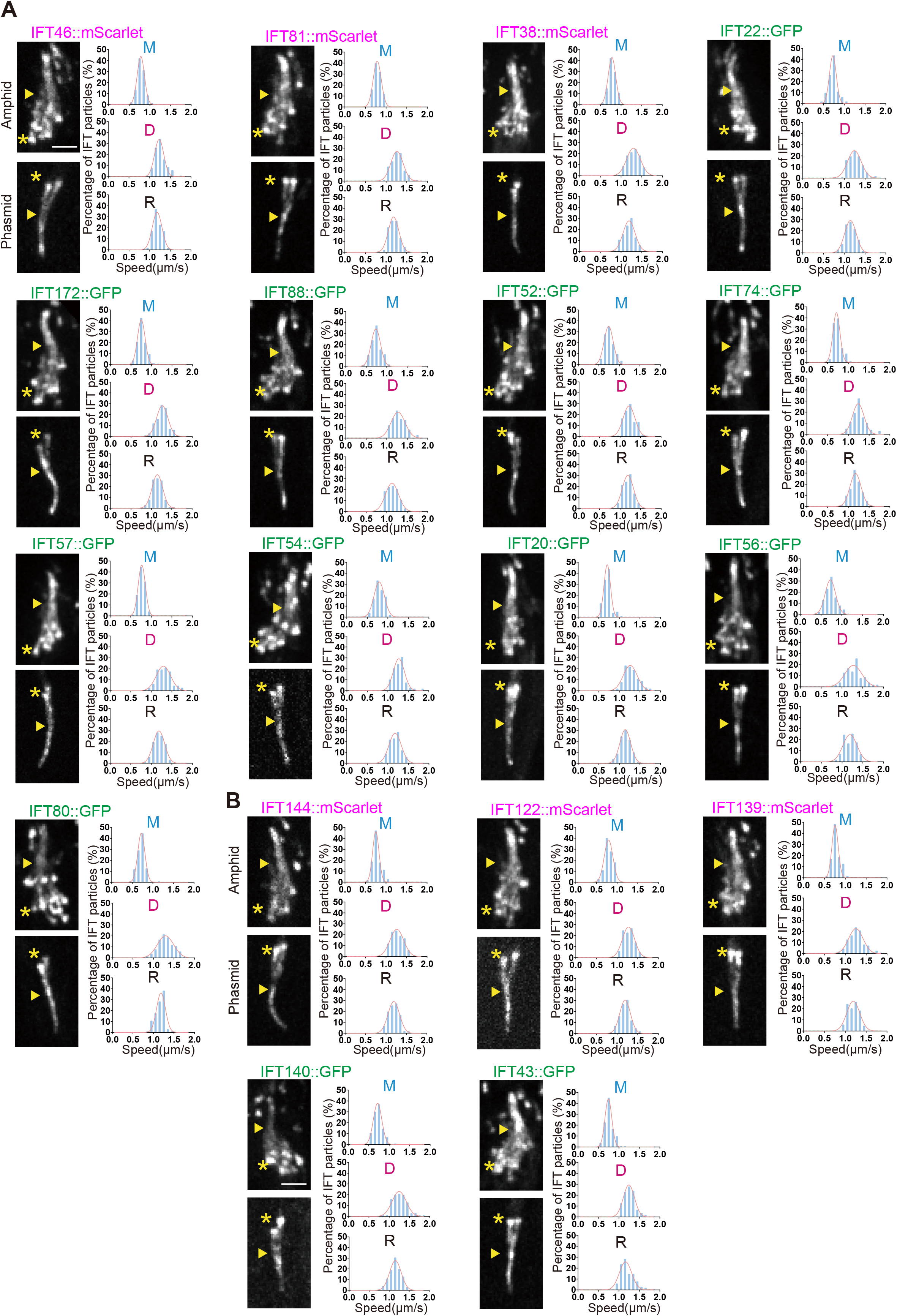
Localization and transport dynamics of endogenous fluorescent knock-in strains for *C. elegans* IFT-particle subunits. **(A)** Representative ciliary images and velocity distributions for endogenous fluorescent knock-in strains of IFT-B subunits, corresponding to the velocity summary shown in Fig. 1A. Left of each strain, representative amphid and phasmid cilia. Asterisks mark the ciliary base/transition zone, and arrowheads mark the junction between the middle and distal segments. Scale bar, 2.5 μm. Right of each strain, velocity distributions and Gaussian fits for IFT transport in the middle segment, distal segment, and retrograde direction. **(B)** Representative ciliary images and velocity distributions for endogenous fluorescent knock-in strains of IFT-A subunits, corresponding to the velocity summary shown in Fig. 1A, displayed as in (A).

**Figure S2.**
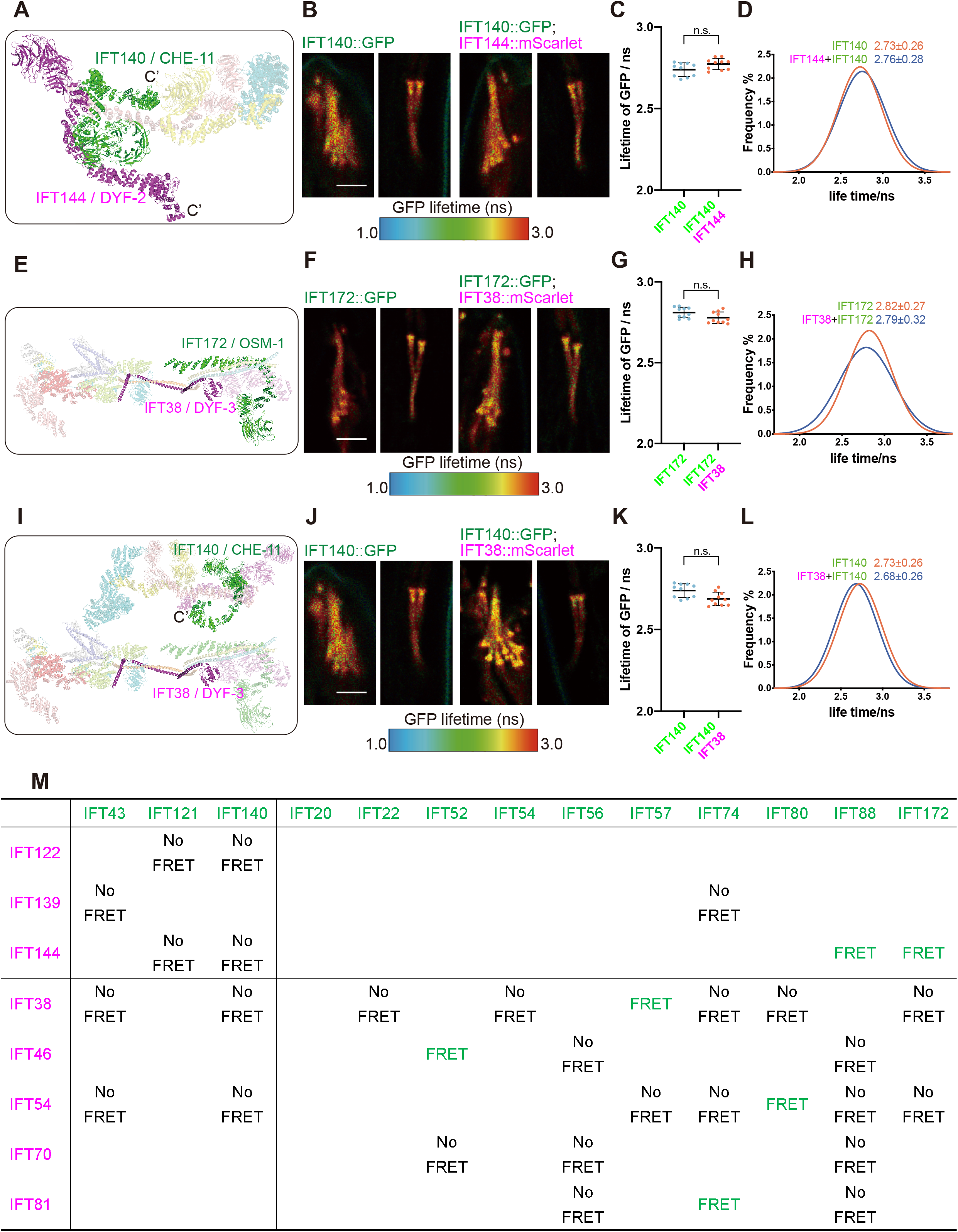
Selected IFT subunit pairs do not show detectable FLIM-FRET under the conditions tested. (A, E, and **I)** Structural models of the IFT complex showing the relative positions of the indicated subunit pairs, including IFT140/CHE-11 and IFT144/DYF-2 (A), IFT172/OSM-1 and IFT38/DYF-3 (E), IFT140/CHE-11 and IFT38/DYF-3 (I). C′ denotes the C-terminal end. **(B, F, and J)** Representative FLIM images of cilia expressing the indicated GFP-tagged donor (green) alone or together with the corresponding mScarlet-tagged acceptor (magenta). GFP fluorescence lifetime is displayed as a pseudocolor map from 1.0 to 3.0 ns. Scale bars, 2.5 μm. **(C, G, and K)** Quantification of GFP lifetimes from the corresponding FLIM images. Each dot represents the apparent mean GFP lifetime of one cilium, calculated by fitting the fluorescence decay curve of that cilium with a mono-exponential decay model. N = 10 cilia from 10 animals per genotype. Bars show mean ± s.d. n.s. not significant, two-tailed Student’s t-test. **(D, H, and L)** GFP fluorescence lifetime dis tributions and Gaussian fits for the corresponding donor-only and donor–acceptor strains. Curves show distributions from the same 10 ciliary regions of interest per genotype. Values in parentheses indicate the mean ± s.d. of the cilium-level apparent mean lifetimes. **(M)** Summary matrix of FLIM-FRET analysis across endogenous GFP- and mScarlet-tagged IFT subunit pairs. Green indicates pairs with detectable FRET, and black indicates pairs with no detectable FRET under the conditions analyzed.

**Figure S3.**
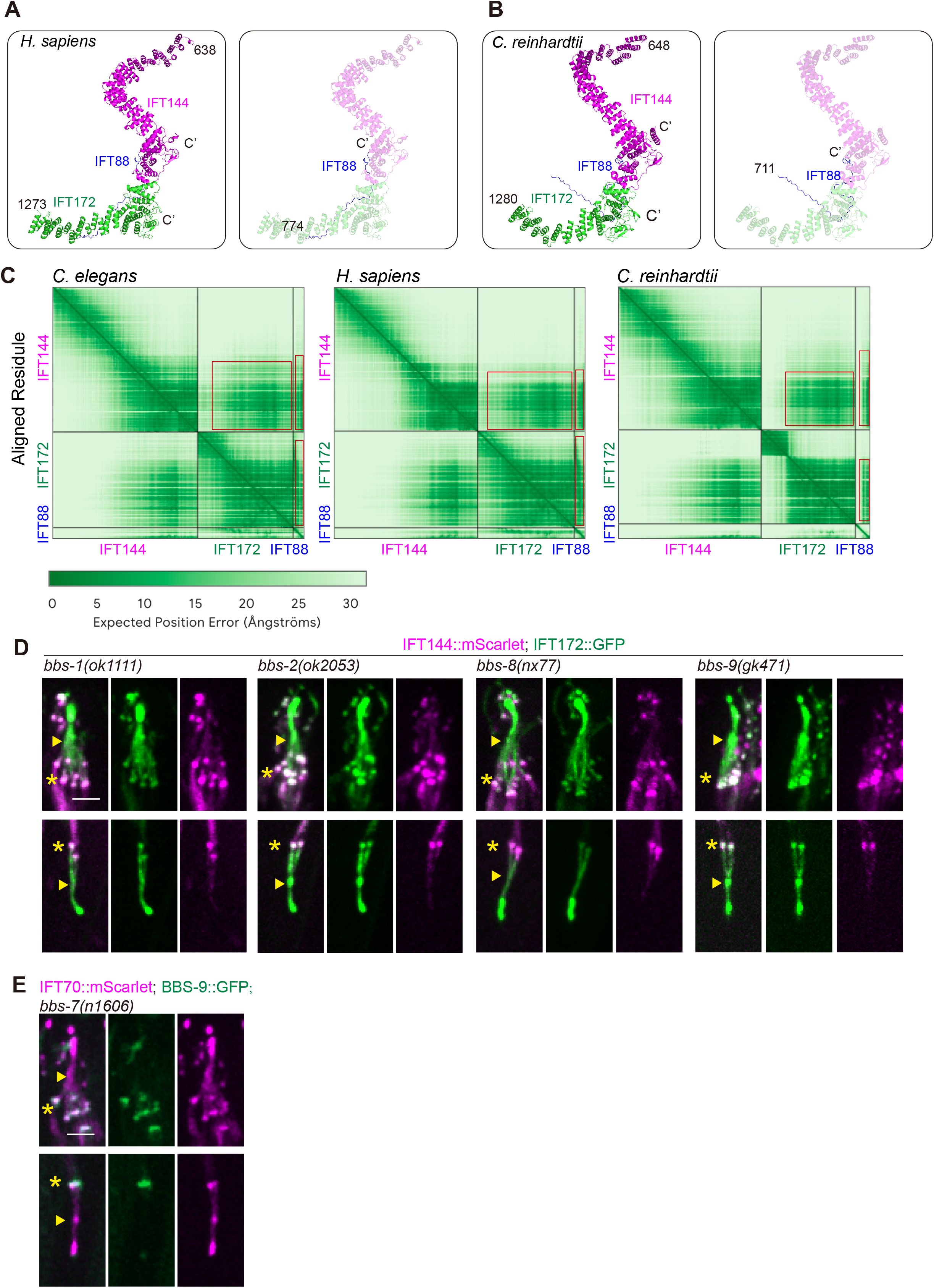
AlphaFold3 predictions of IFT-A and IFT-B relationships across species and phenotypes of *bbs* mutants in *C. elegans*. **(A)** AlphaFold3-predicted model of C-terminal regions of human IFT144, IFT172 and IFT88. **(B)** AlphaFold3-predicted model of C-terminal regions of *Chlamydomonas reinhardtii* IFT144, IFT172 and IFT88. **(C)** PAE plots of AlphaFold3-predicted models of Fig. 3B and Fig. S3A-B. Red boxes highlight possible interactions between IFT144, IFT172 and IFT88. **(D)** Representative ciliary images of endogenous IFT144/DYF-2::mScarlet and IFT172/OSM-1::GFP in *bbs-1(ok1111)*, *bbs-2(ok2053)*, *bbs-8(nx77*), *bbs-9(gk471)* mutant animals. Asterisks mark the ciliary base/transition zone, and arrowheads mark the junction between the middle and distal segments. Scale bar, 2.5 μm. **(E)** Representative ciliary images of endogenous IFT70/DYF-1::mScarlet and BBS-9::GFP in *bbs-7(n1606)* mutant animals. Asterisks mark the ciliary base/transition zone, and arrowheads mark the junction between the middle and distal segments. Scale bar, 2.5 μm.

**Figure S4.**
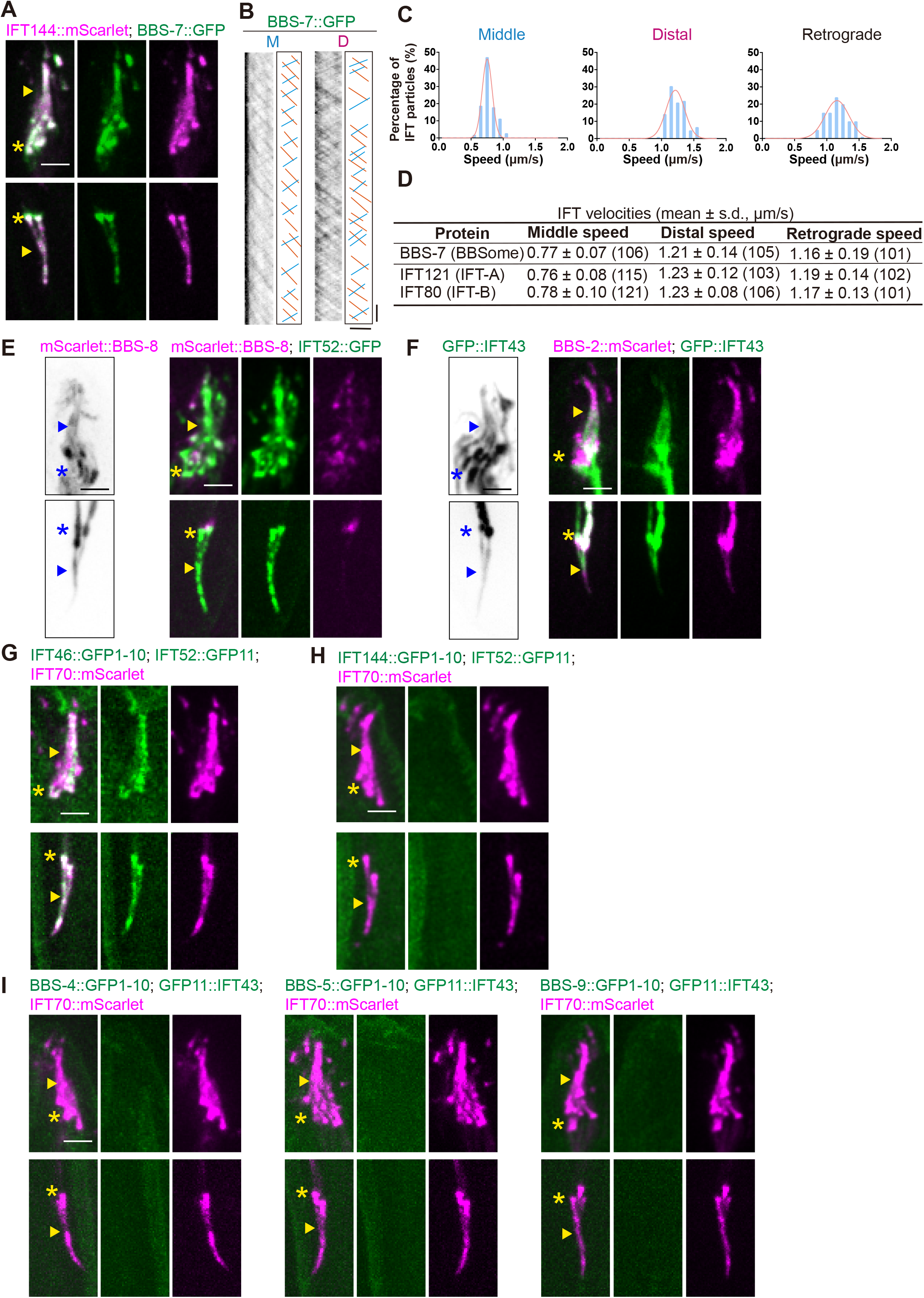
BBSome transport dynamics and controls for BBSome–IFT proximity assays. (A) Representative ciliary images of endogenous BBS-7::GFP together with IFT144/DYF-2::mScarlet in *C. elegans* sensory cilia. Asterisks mark the ciliary base/transition zone, and arrowheads mark the junction between the middle and distal segments. Scale bar, 2.5 μm. **(B)** Representative kymographs showing BBS-7::GFP transport. Orange lines indicate anterograde tracks, and blue lines indicate retrograde tracks. Horizontal scale bar, 2.5 μm; vertical scale bar, 2.5 s. **(C)** Velocity distributions and Gaussian fits for BBS-7::GFP transport in the middle segment, distal segment, and retrograde direction. **(D)** Summary of transport velocities for BBS-7::GFP, the representative IFT-A subunit IFT121/IFTA-1, and the representative IFT-B subunit IFT80/CHE-2. Data are mean ± s.d.; numbers in parentheses indicate the number of transport events analyzed. Data are from at least 10 independent phasmid cilia of *C. elegans*. **(E)** Representative ciliary images of mScarlet::BBS-8 expressed alone or co-expressed with IFT52/OSM-6::GFP. Asterisks and arrowheads are as in (A). Scale bar, 2.5 μm. **(F)** Representative ciliary images of GFP::IFT-43 expressed alone or co-expressed with BBS-2::mScarlet. Asterisks and arrowheads are as in (A). Scale bar, 2.5 μm.**(G)** Positive control for split-GFP complementation between IFT46/DYF-6 and IFT52/OSM-6. IFT70/DYF-1::mScarlet was used as a ciliary IFT marker. Asterisks and arrowheads are as in (A). Scale bar, 2.5 μm. **(H)** Structure-guided negative control for split-GFP complementation between the spatially separated IFT-A subunit IFT144/DYF-2 and IFT-B subunit IFT52/OSM-6. IFT70/DYF-1::mScarlet was used as a ciliary IFT marker. Asterisks and arrowheads are as in (A). Scale bar, 2.5 μm. **(I)** Split-GFP complementation tests between IFT43 and the BBSome subunits BBS-4, BBS-5, or BBS-9. No detectable split-GFP signal was observed for these pairs under the same imaging conditions. IFT70/DYF-1::mScarlet was used as a ciliary IFT marker. Asterisks and arrowheads are as in (A). Scale bar, 2.5 μm.

**Figure S5.**
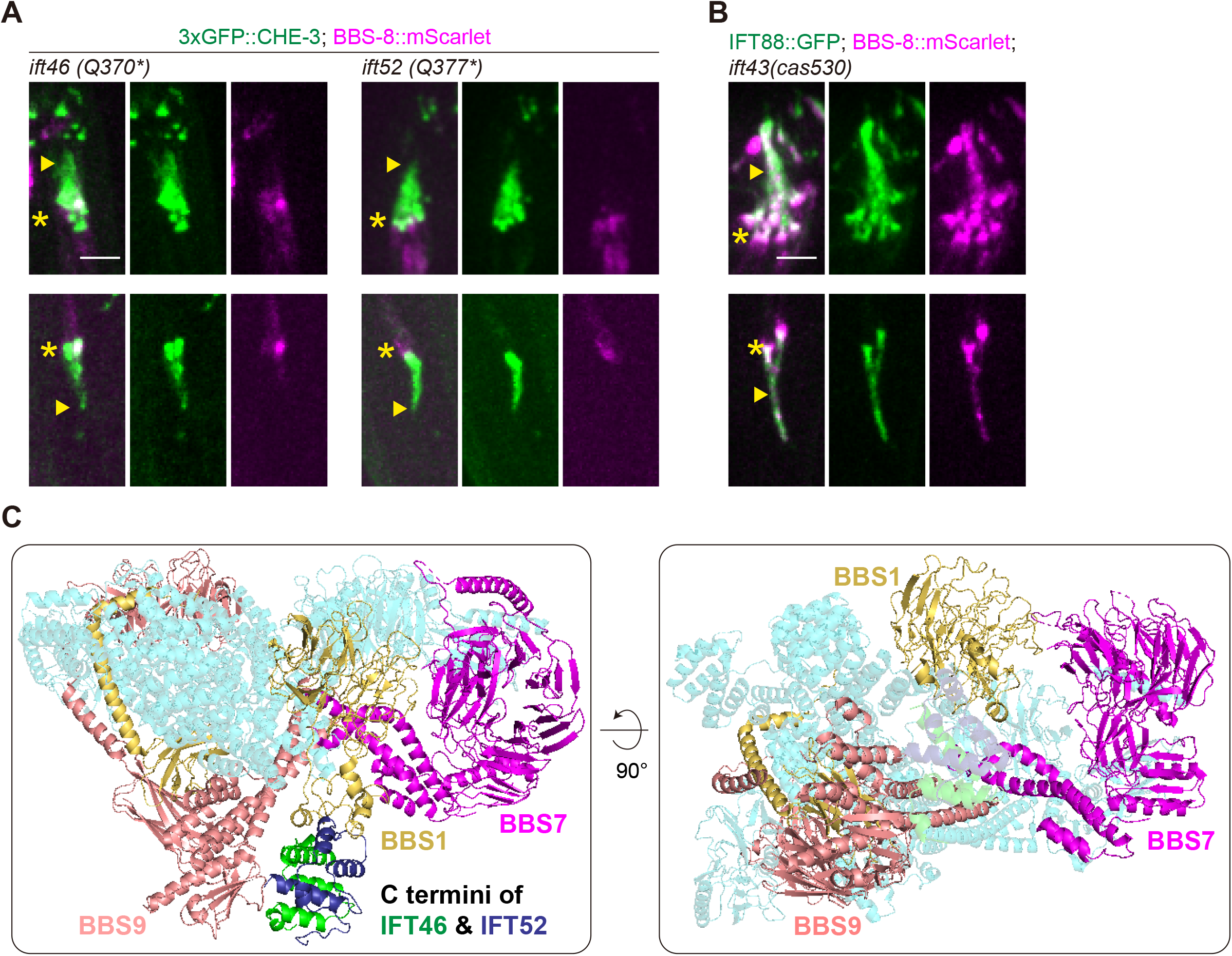
C-terminal IFT46–IFT52 truncation mutants show reduced BBS-8 ciliary entry. (A) Representative ciliary images of BBS-8::mScarlet in *ift46(Q370*)* and *ift52(Q377*)* animals, which carry C-terminal truncations of IFT46/DYF-6 and IFT52/OSM-6, respectively. 3xGFP::CHE-3 was used as a ciliary marker. Asterisks mark the ciliary base/transition zone, and arrowheads indicate the distal end of the remaining middle segment. Scale bar, 2.5 μm. **(B)** Representative ciliary images of BBS-8::mScarlet in *ift43(cas530)* mutant animals. IFT88/OSM-5::GFP was used as a ciliary IFT marker. Asterisks and arrowheads are as in (A). Scale bar, 2.5 μm. **(C)** AlphaFold3-based model of the *C. elegans* BBSome with the C-terminal regions of the IFT-B subunits IFT46/DYF-6 and IFT52/OSM-6. Shown are IFT46/DYF-6 residues 368–446 and IFT52/OSM-6 residues 369–427. BBS-1, BBS-7, and BBS-9 are highlighted. The predicted *C. elegans* BBSome model was aligned to the reported bovine BBSome structure (6VNW). The right panel shows the same model after 90° rotation.

**Figure S6.**
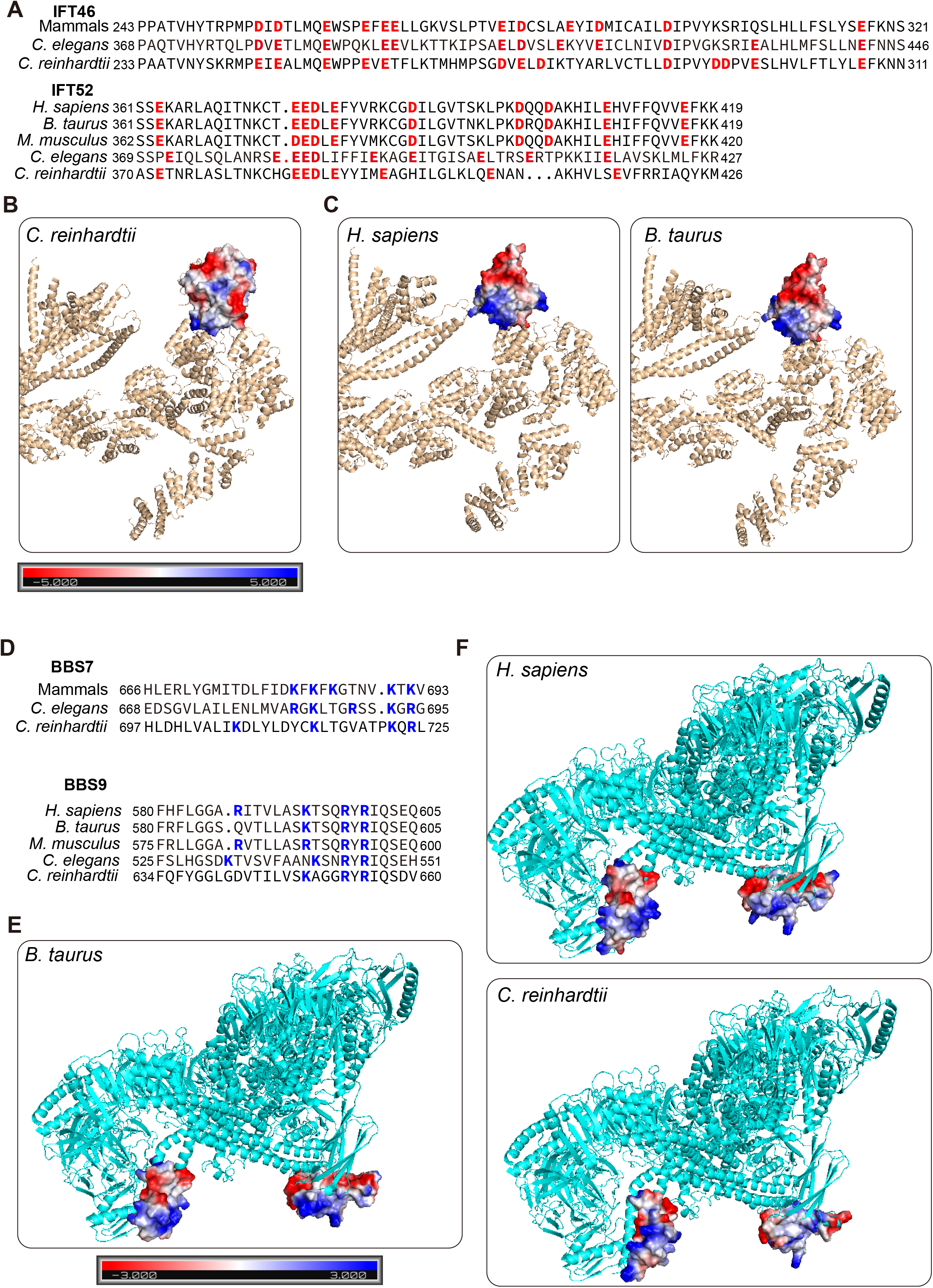
Electrostatic features of the candidate BBSome–IFT-B interface are conserved across species. (A) Sequence alignment of the IFT46 and IFT52 C-terminal regions from human, mouse, bovine,*C. elegans* and *C. reinhardtii*. Acidic residues are highlighted in red. Homologous sequence alignment was performed by CLUSTALW. **(B)** Surface electrostatic potential of the *C. reinhardtii* IFT46–IFT52 C-terminal module, analyzed from the reported IFT-B structural model (8BD7). Surface electrostatic potential was calculated using the APBS plugin in PyMOL and displayed on a scale from −5 to +5 kBT/e. **(C)** Surface electrostatic potential of the human and bovine IFT46–IFT52 C-terminal modules. The C-terminal regions of IFT46 and IFT52 were modeled using AlphaFold3 and aligned to the reported IFT-B structure (8BD7) before electrostatic analysis. **(D)** Sequence alignment of BBS7 and BBS9 regions from human, mouse, bovine, *C. elegans* and *C. reinhardtii*. Basic residues are highlighted in blue. Homologous sequence alignment was performed by CLUSTALW. **(E)** Surface electrostatic potential of the candidate BBS7 and BBS9 region in the reported bovine BBSome structure (6VNW). Surface electrostatic potential was calculated using the APBS plugin in PyMOL and displayed on a scale from −3 to +3 kBT/e. **(F)** Surface electrostatic potential of the corresponding candidate BBS7 and BBS9 regions in human and C. reinhardtii models. The structures were predicted using AlphaFold3, aligned to the reported BBSome structure (6VNW) and analyzed as in (E).

**Figure S7.**
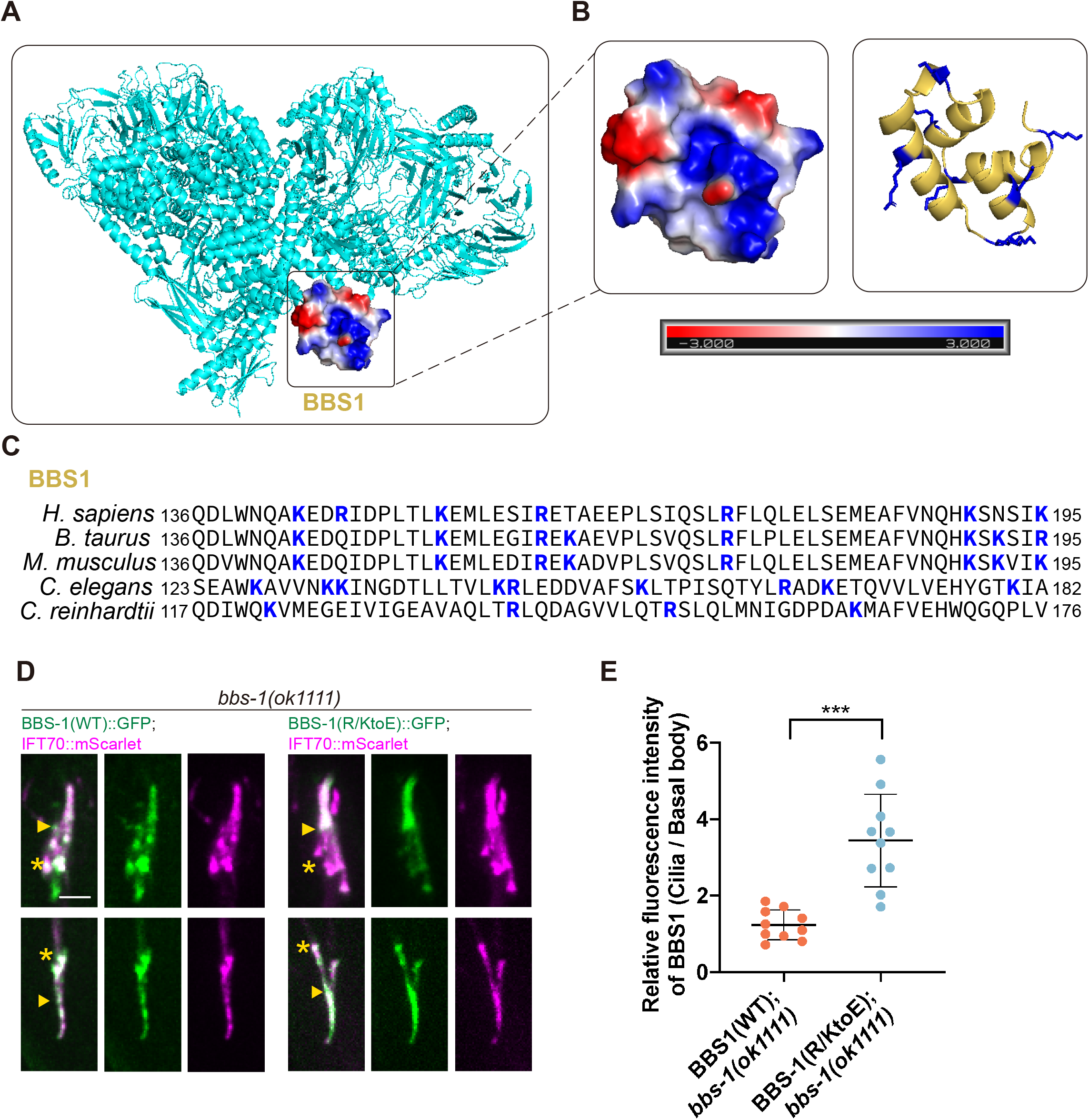
Charge reversal of the candidate BBS-1 basic surface does not prevent BBS-1 ciliary entry. (A) Surface electrostatic potential of candidate BBSome regions containing BBS1. The *C. elegans* BBSome model was predicted using AlphaFold3 and aligned to the reported bovine BBSome structure (6VNW). Boxed regions indicate candidate electropositive surfaces. Surface electrostatic potential was calculated using the APBS plugin in PyMOL and displayed on a scale from −3 to +3 kBT/e. **(B)** Close-up view of the candidate BBS-1 surface shown in (A). Left, surface electrostatic potential; right, structural model with basic side chains highlighted in blue. **(C)** Sequence alignment of the BBS-1 region analyzed in (A and B) from human, bovine, mouse, *C. elegans*, and *C. reinhardtii*. Basic residues are highlighted in blue. Homologous sequence alignment was performed by CLUSTALW. **(D)** Representative ciliary images of *bbs-1(ok1111)* animals expressing IFT70::mScarlet together with either wild-type BBS-1::GFP or the charge-reversal BBS-1(R/KtoE)::GFP variant. BBS-1(R/KtoE) denotes a charge-reversal variant in which all the basic residues highlighted in blue of *C. elegans* sequences in (C) were substituted with glutamate. Asterisks mark the ciliary base/transition zone, and arrowheads mark the junction between the middle and distal segments. Scale bar, 2.5 μm.**(E)** Quantification of BBS-1::GFP ciliary enrichment in the wild-type and charge-reversal conditions shown in (D). Ciliary enrichment was calculated as the fluorescence intensity in the ciliary shaft divided by the fluorescence intensity at the basal body. Ten phasmid cilia were analyzed per condition. Data are mean ± s.d. P values were calculated using one-way ANOVA followed by Tukey’s multiple-comparisons test. ***P < 0.001.

